# Single-molecule mechanical unfolding kinetics of unmodified *Saccharomyces cerevisiae* tRNA^Phe^: a hint to the tRNA chaperone-tRNA interaction mechanism

**DOI:** 10.1101/2021.05.03.442431

**Authors:** Wenzhao Liu, Luyi Feng, Wenpeng Zhu, Zhenyu Zou, Ran Chen, Jie Zhou, Wei Xie, Hu Chen, Zhensheng Zhong, Jie Ma

**Affiliations:** School of Physics, Sun Yat-sen University, Guangzhou 510275, Guangdong, China; State Key Laboratory of Optoelectronic Materials and Technologies, Sun Yat-sen University, Guangzhou 510275, Guangdong, China; State Key Laboratory for Biocontrol, School of Life Sciences, Sun Yat-sen University, Guangzhou 510060, Guangdong, China; Research Institute for Biomimetics and Soft Matter, Fujian Provincial Key Lab for Soft Functional Materials Research, Department of Physics, Xiamen University, Xiamen 361005, Fujian, China

## Abstract

The biological activity of tRNA is closely related to its mechanical folding properties. Although previous studies focused on the folding and unfolding mechanism of tRNA, its kinetics are largely unknown. In this study, combining optical tweezers and molecule dynamics simulations, we characterized the mechanical folding and unfolding processes of a single unmodified *Saccharomyces cerevisiae* tRNA^phe^. We identified the intermediates and pathways for tRNA mechanical folding and unfolding in the presence of Mg^2+^, discovering that the folding/unfolding kinetics of D stem-loop and T stem-loop but not the anti-codon stem-loop significantly affected by their upstream and downstream structures. The cooperative unfolding of the tRNA in the presence of Mg^2+^ lead to a large hysteresis between the folding and unfolding pathway, and such hysteresis and unfolding cooperativity are significantly reduced by lowering the Mg^2+^ concentration or mutating the nucleotides forming the ‘elbow’ structure. Moreover, both steered molecular dynamics simulation and optical tweezers experiment results support that, formation of tertiary interactions in the elbow region increases energy barriers of the mechanical unfolding pathway, including those in between intermediates, and determines the overall unfolding cooperativity. Our studies may shed light on the detailed tRNA chaperone mechanism of TruB and TrmA.

## INTRODUCTION

The folding state and folding kinetics affect many processes in the life cycle of a tRNA molecule. In the tRNA maturation and degradation processes in eukaryotic cells, proteins or complexes such as Lupus autoantigen, RNase P, exportin-t, tRNA nucleotidyltransferase and xrn-1 interact with pre-tRNA or tRNA substrates by virtue of their structural characteristics (1–5). Moreover, tRNAs may change their folding state during some essential biological processes. For example, tRNA is subject to tension and is distorted by the ribosome during its translocation (6). Also importantly, the tRNA folding states *in vivo* could be perturbed kinetically by RNA helicases, which is in sharp contrast with previous tRNA folding studies under an equilibrium condition (7,8) and hence demands a better understanding of tRNA folding/unfolding kinetics under force.

Recently, it is discovered that some RNA modification enzymes such as *Escherichia coli* TruB and TrmA bind and partially unfold tRNAs transiently as tRNA chaperones (9,10). Both tRNA chaperones specifically bind to the elbow region of tRNA, and such ability is suggested to related to the chaperone activity (11). However, the elbow region may not form when the tRNA substrates are kinetically trapped in partially folded or misfolded conformations (12). Therefore, how the tRNA chaperones resolved these kinetically trapped substrates without ATP hydrolysis is still not clear. Unraveling the detailed mechanical folding and unfolding kinetics of native tRNA and tRNA intermediates may help understand the mechanism of these tRNA-associated biological functions.

Interestingly, despite the great success achieved in tRNA folding research (13–19), the folding and unfolding kinetics of tRNAs, especially kinetics of tRNA intermediates remain largely unknown. Although previously single-molecule fluorescence resonance energy transfer (smFRET) technique has been successfully used to reveal the detailed folding kinetics of human mitochondrial tRNA^lys^ (which is unstable at near physiological conditions and have alternative states), the tRNAs which can be studied in this way at near physiological conditions are rather limited (20,21). In contrast, for most canonical tRNAs, the native L-shape folded status is predominated under physiological conditions and is rather stable. For these majorities of tRNAs, the detailed folding and unfolding mechanisms in physiological conditions still remain elusive. A proof-of-concept study mechanically unfolded individual canonical tRNAs using α-hemolysin nanopore (22). However, it is difficult to unravel the folding and unfolding kinetics of tRNA intermediates from the nanopore translocation signals.

On the other hand, optical tweezers based single-molecule force spectroscopy enables direct evaluation of the mechanical stability of RNA structures at near physiological conditions, and extraction of force-dependent kinetics and free energy landscapes of the entire folding/unfolding pathways (23,24). In this study, using *Saccharomyces cerevisiae* cytosolic tRNA^phe^ transcript (Figure. 1A) as a model, we studied the mechanical folding/unfolding properties of type I tRNA at different Na^+^ and Mg^2+^ concentrations. Two types of optical-trapping experiments were carried out here: one is the pulling experiment based on single-trap assay (Figure. 1B, top panel) which is more convenient in forming tethers and acquiring data; the other is the force-clamp experiment based on dual-trap assay which allows higher precision (Figure. 1B, bottom panel) and is less perturbed by the stage drifting. To gain more insight into tRNA folding/unfolding at the atomic level, we also performed equivalent steered molecular dynamics (SMD) simulations. Together, these studies unraveled the detailed unfolding and folding mechanisms of tRNA under different buffer conditions, which clearly exhibit the coupling between secondary and tertiary structures in tRNA folding and unfolding. Our study also demonstrated that the formation of the tertiary interactions in the elbow region increases energy barriers in the tRNA unfolding pathway, which suggests that tRNA chaperone resolved the kinetically trapped misfolded and partially folded tRNA by disrupting these interactions even when the elbow structure is not fully formed.

**Figure 1.**
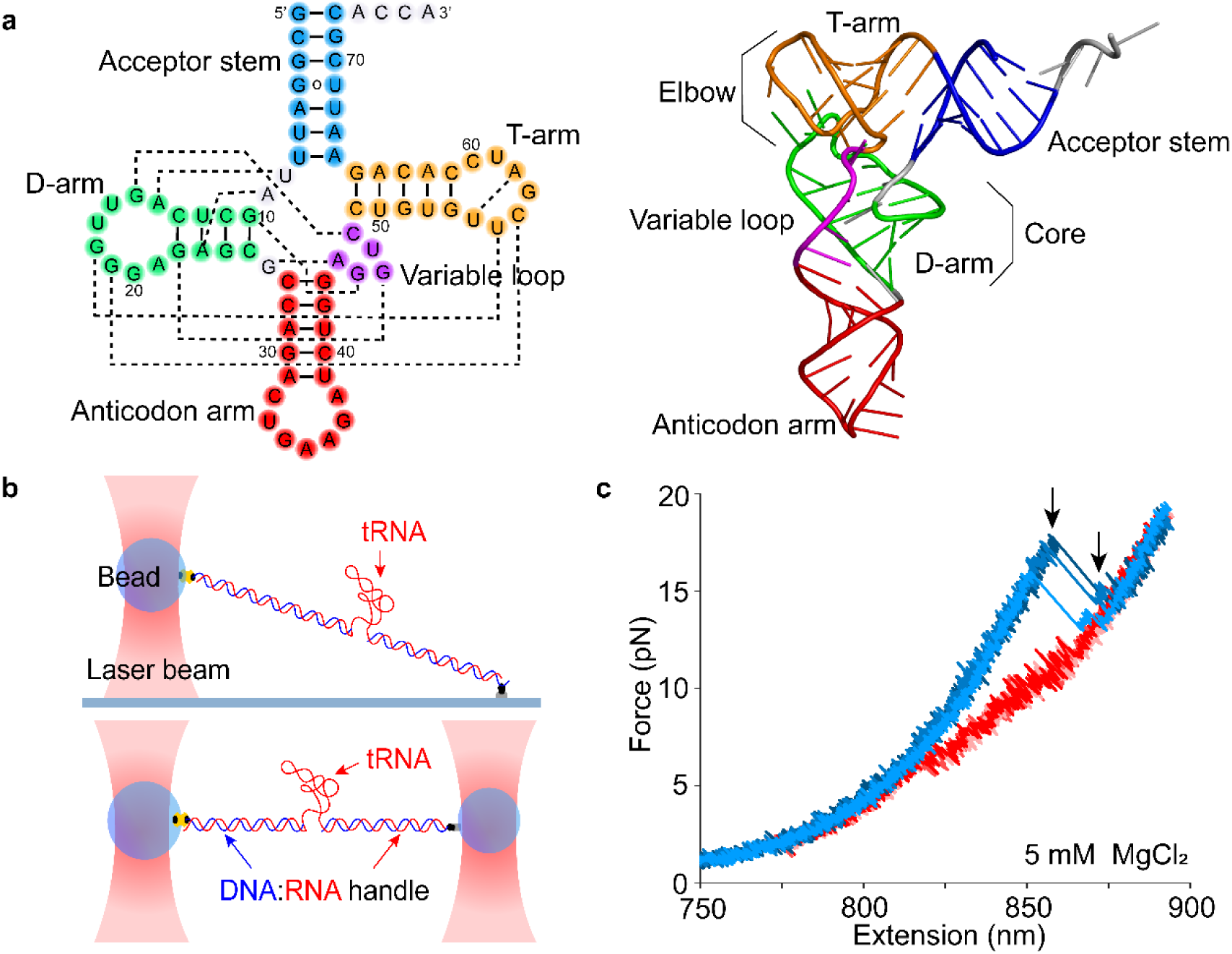
Scheme of the experiments. (A) The cloverleaf secondary and L-shaped tertiary structure of the cytosolic tRNA^phe^ from *S. cerevisiae*. Protein Data Bank entry (PDB): 1EHZ; (B) schematic of the optical-tweezers based single-molecule folding/unfolding experiments. Upper panel: In the pulling experiment, tRNA molecules were tethered between streptavidin-coated polystyrene bead and coverglass surface. Lower panel: In the force-clamp experiment, tRNA molecules were tethered between streptavidin-coated polystyrene bead and anti-Digoxigenin-coated polystyrene bead. (C) Representative FECs of unfolding (blue) and refolding (red) of tRNA from three different tethers during the pulling experiments at 5 mM MgCl_2_. The two unfolding ruptures were indicated by black arrows.

## MATERIAL AND METHODS

### Sample preparation

Single-molecule constructs of the wildtype and mutant yeast tRNA^phe^ samples were generated as described previously (25). In brief, the recombinant plasmids were constructed by inserting chemically synthesized DNAs into the pUC19 vector. The 2725-nt RNA molecules were synthesized by polymerase chain reaction (PCR) amplification of the plasmids followed by *in vitro* transcription using the T7 RNA polymerase (Promega, Fitchburg, WI, USA). The RNAs contain a 1206-nt upstream sequence, a 2-nt linker (CC), the 76-nt tRNA sequences with the CCA tail, a downstream 1-nt linker (T), and a 1440-nt downstream sequence (Supplementary Figure S1). The RNAs were annealed with complementary strands of PCR-generated 1206-bp and 1440-bp dsDNAs to generate RNA/DNA hybrid upstream and downstream handles, respectively. The upstream handle was labeled at the 3’ end of the DNA strand by introducing biotin-16-dUTP (Biotium, Fremont, CA, USA) using the T4 DNA polymerase (New England Biolabs, Beverly, MA, USA). The downstream handle was labeled at the 5’ end of the DNA strand by digoxigenin-labeled primer (Sangon, Shanghai, China) during PCR.

Single-molecule constructs of the individual anti-codon stem-loop (ASL), D stem-loop (DSL) and T stem-loop (TSL) as well as the truncated tRNA samples DAV (the 10^th^ to 48^th^ nucleotides of the tRNA) and AVT (the 27^th^ to 65^th^ nucleotides of the tRNA) were generated using another strategy described by Block *et al* (26). Briefly, RNAs containing the targeted sequences and the upstream and downstream 1-nt flanking sequences in the wildtype tRNA^phe^, and the upstream 30-nt and downstream 30-nt ‘sticky’ sequences, were synthesized by *in-vitro* transcription using the T7 RNA polymerase (Promega) and synthetic DNA templates (Sangon). Two dsDNA handles were generated by PCR using the pUC19 plasmid as the template. The 1195-bp upstream handle contains an abasic site and a 30-nt 5’ overhang introducing by an autosticky primer (27), and a digoxigenin label at the opposite 5’ end introducing by the other primer. The 1409-bp downstream handle with a 30-nt 3’ overhang was generated by PCR using a primer with 5’-phosphorylation and three phosphorothioate bonds and another with 5’-biotin modification, followed by 1-minute lambda exonuclease (New England Biolabs) digestion (28,29). The RNA was annealed to the handles with the ratio of 1:1:1 in a buffer containing 72% formamide, 33 mM piperazine-1,4-bisethanesulfonic acid (PIPES) (Sigma-Aldrich, St. Louis, MO, USA), 300 mM NaCl, 0.8 mM EDTA, pH 7.0. During the annealing process, the temperature was first held at 85 °C for 10 min, then kept at 62 °C for 90 min, and held at 52 °C for 90 min, cooled to 4 °C at a rate of −1 °C/min. All the synthetic DNA sequences used in this study are listed in the Supplementary Table 12.

### Optical trapping and Data acquisition

Single-molecule pulling experiments were performed using home-built single-trap optical tweezers described previously (30). Briefly, a linearly polarized TEM_00_-mode 1064-nm laser beam (Changchun New Industries Optoelectronics Technology, Changchun, China) was coupled to a polarization-maintaining single-mode fiber (OZ Optics, Ottawa, Canada). Then, after two beam expanders, the beam was focused at the sample plane by a 100X, 1.45-NA, oil immersion objective mounted on an inverted microscope (Ti-U microscope, Nikon, Tokyo, Japan). The displacements of the trapped beads from the trap center were measured by a position-sensitive detector (First Sensor, Berlin, Germany) and later converted into a force signal. Analog voltage signal was digitized at 10 kHz for each channel using a multiplexed analog-to-digital conversion PCI-E board (National Instruments, Austin, TX) and averaged to 1 kHz. Calibration and data conversion methods were adapted from those used by Wang *et al* (31). The samples were tethered to a 0.8-μm streptavidin-coated polystyrene bead (Spherotech, Lake Forest, IL) at one end through biotin-streptavidin interactions and to the anti-digoxigenin (Roche, Basel, Switzerland) coated coverglass surface at the other end through digoxigenin-antibody interactions. The polystyrene bead was optically trapped and the coverglass was driven by a servo-controlled 3D piezoelectric stage (Physik Instrumente, Karlsruhe, Germany). The stage moved at a constant speed while the bead displacement from the trap center was monitored by the PSD. The laser power was held constant during the whole measurement. Stretching and relaxing cycles were performed at a piezo moving speed of 50 nm/s, resulting in an effective force-loading rate about 4.7 pN/s in the force range of 5-20 pN. Each tether was pulled no more than 5 times.

Force-clamp experiments were performed on a commercial dual-trap optical tweezers (C-Trap, LUMICKS, Amsterdam, Netherlands). Briefly, a ~10 W 1064-nm fiber laser was split by the polarization beam splitter cube into two independent beams, which were steered by two piezo-driven deflectors, one for the coarse-positioning of trap 2 and other for the accurate-positioning of trap 1. The beams were recombined and expanded before entering a water-immersion objective. A linear stage was used to rapidly switch the optical traps between five channels of the flow cell, allowing *in situ* construction of the dumbbell constructs. Channels 1-3 of the flow cell were separated by a laminar flow, which contained 0.8-μm streptavidin-coated polystyrene beads (Spherotech), the buffer, 0.5-μm anti-Digoxigenin coated polystyrene beads (a gift from Assistant Professor Bo Sun at ShanghaiTech University), respectively. The streptavidin-coated polystyrene beads were incubated with single-molecule constructs for about 30 minutes at room temperature before injecting to Channel 1. The intensities of the two beams were modulated to achieve approximately equal stiffness, with a typical value of 0.1-0.2 pN/nm. A single streptavidin-coated bead and a single anti-Digoxigenin-coated bead were caught in each trap in channel 1 and channel 3, and then quickly moved to channel 2. The dumbbell construct was formed by moving one trap close to the other and incubated for 10-30 s, and the formation of tether was indicated by changes in the force signals. Several FECs were obtained prior to force-clamp experiments in the buffer channel. Tethers with obviously shorter contour lengths (compared to the theoretical WLC curve), which tended to be multiple tethers, were discarded. All the force-clamp data were collected at 78.125 kHz using the Bluelake software provided by LUMICKS and further analyzed using custom Matlab programs.

For both pulling and force-clamp experiments, the buffer conditions are 10 mM PIPES, pH 7.0, 0.5 mM EDTA, 0.2 U/μL RNasin plus RNase Inhibitor (Promega), 4 mM protocatechuic acid (PCA) (Sigma-Aldrich) and 100 nM protocatechuate 3,4-dioxygenase (PCD) (Sigma-Aldrich) at indicated NaCl concentrations or 10 mM PIPES, pH 7.0, 50 mM KCl, 0.2 U/μL RNasin plus RNase Inhibitor, 4 mM PCA and 100 nM PCD at indicated MgCl_2_ concentrations. All the experiments were performed at a temperature (24 ± 1°C) and humidity (50 ± 5%) controlled room. Data analysis methods for optical tweezers experiments were described in detail in the Supplementary Text.

### Molecular Dynamic (MD) simulation

Coarse-grained and all-atom simulations based on the GROMACS package (version 2018.4) (32) were performed to investigate the unfolding dynamics of tRNAs and the stability of the tertiary structures, respectively. In the all-atom simulations, the AMBER-14SB force field (33–35) was employed for the bonded and non-bonded interactions between atoms of tRNA molecules in the simulations. The canonical L-shaped folded tRNA (PDB ID: 1EHZ) or the corresponding mutant tRNA was initially solvated in the water box with the same structure and orientation. For enhanced computational efficiency, water molecules were represented by the TIP3P model (36). For the cases with cations, Mg^2+^ or Na^+^ ions were added into the simulation system to neutralize the negative charge of the tRNAs. To simulate the tRNAs without cations, a uniform charge distribution was added to the background to compensate the net charge of the system. The fast smooth particle-mesh Ewald (37) was used to calculate the long-range electrostatic interactions. The system was modeled as an NPT ensemble, with periodic boundary conditions in all directions, under constant pressure *P* (1 atm) and constant temperature *T* (300 K). The simulation system containing 66,000 water molecules covered the dimensions of 15 nm × 15 nm ×15 nm, which was large enough to prevent the tRNA molecule from interacting with the periodic images. The time step was fixed at 2 fs. After 20 ns of equilibration, steered molecular dynamics (SMD) simulations (38,39) were performed to separate the base pairs in the elbow structure or the AS structure. The separation rate was 0.05 nm/ns. To evaluate the stability of the tertiary structures, we used thermodynamic integration technique to calculate the free binding energy of the base pairs in the elbow structure and the AS structure as a function of the separation distance (25).

In the Coarse-grained simulations, the Martini CG force field (40) was employed to reduce the computational costs and perform large time-space scale simulations parallel to the experiments of tRNA unfolding. The elastic networks in Martini CG model were modified to capture the transformation of the secondary and tertiary structures. The CG structures of the canonical L-shaped folded tRNA and the elbow-disrupted folded tRNA were constructed based on the equilibrated structures in the all-atom simulations. The tRNA molecule was solvated in the CG water box containing 165,000 water molecules and having dimensions of 15 nm × 15 nm ×70 nm, which was necessary to model the entire dynamics of tRNA unfolding at the time scale up to microseconds. Mg^2+^ ions were added into the system to neutralize the negative charge of tRNAs, while the uniform charge distribution was added to the background to model the case without cations. The reaction-field (41) was used to calculate the long-range electrostatic interactions. The system with the periodic boundary conditions in all directions was simulated in an NPT ensemble under constant pressure *P* (1 atm) and constant temperature *T* (300 K). The time step was fixed at 5 fs. After the initial equilibration, the SMD simulations of tRNA unfolding were performed under the loading rates of 0.2, 0.5, 1, 2, 5, 10 nm/ns. The spans for the disruptions of secondary structures were identified between the critical extensions when the first and the last base pair broke for each stem-loop.

## RESULTS

### Wildtype unmodified yeast tRNA^phe^ unfolds cooperatively in two steps and refolds in multiple steps in the presence of magnesium ions

Firstly, we used the wildtype tRNA in the presence of 5 mM MgCl_2_ for our single-molecule mechanical unfolding experiments. We combined both force-extension curves (FECs) at constant pulling speed and trajectories at constant force to identify states in the unfolding and folding pathways. Experimental details can be found in the Methods section. Typical stretching and relaxation FECs of wildtype tRNA in the presence of 5 mM MgCl_2_ were shown in Figure 1C. As can be seen, a two-step unfolding pathway was observed with one large transition occurring at 16-18 pN, followed by a small hopping transition at 12-14 pN. No more transitions were observed when the force exceeded 20 pN (data not shown). However, when the tRNA refolded, it followed a multiple-step pathway. To further identify these intermediate states, we performed the high-resolution force-clamp experiments at 5 mM MgCl_2_ with dual-trap optical tweezers (Figure 2A).

**Figure 2.**
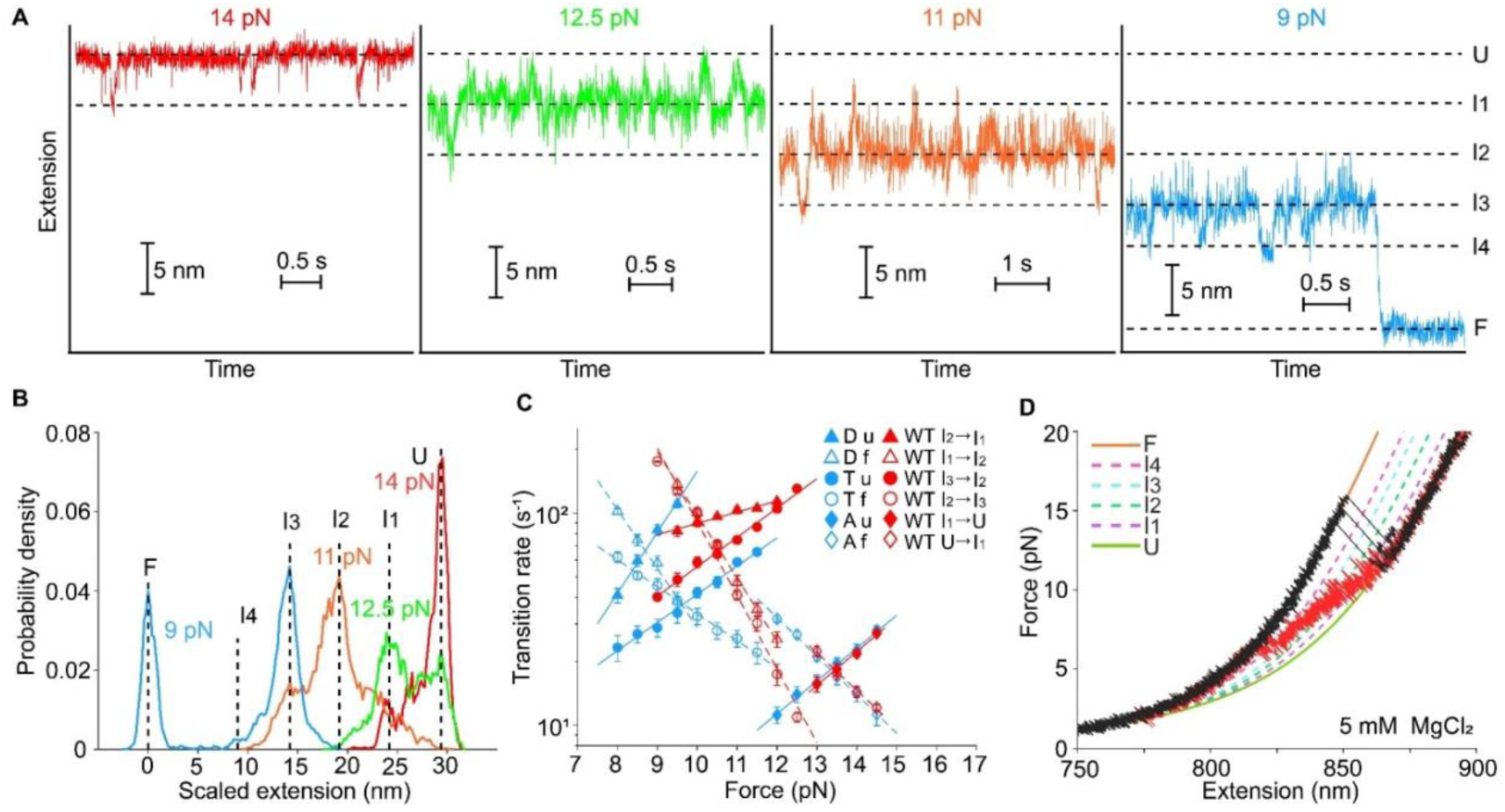
Mechanical unfolding and folding of wildtype tRNA^phe^ at 5 mM MgCl_2_. (A) Typical force-dependent folding trajectories of wildtype tRNA^phe^ at 5 mM MgCl_2_ in the force-clamp experiment. Six distinct states were revealed. (B) Distributions of extension changes from typical traces (~60 s) at different forces showing force-dependent occupancies of the states. (C) Force-dependent kinetics of unfolding/folding at each transition of wildtype tRNA^phe^ as well as that of individual ASL, DSL and TSL during force-clamp experiments. Solid symbols represent the unfolding process and hollow symbols represent the folding process. (D) Representative FECs of unfolding (black) and refolding (red) of tRNA during the pulling experiments at 5 mM magnesium ion. Extensible WLC curves are shown for the six states: ‘F’, (orange); ‘U’, (yellow-green); ‘I_1_’, (purple); ‘I_2_’, (green); ‘I_3_’, (cyan); ‘I_4_’, (pink).

During the force-clamp experiments, the tRNA was first fully unfolded at high force (i.e. 18 pN). After that, the force was reduced step by step (0.5 pN for each step) and the extension was measured while the force was kept constant at each step (Supplementary Figure S6A). The force was stepped down until tRNA was completely folded. In the presence of 5 mM MgCl_2_, transitions among six different extensions were observed, indicating four intermediate states (i.e. ‘I_1_’, ‘I_2_’, ‘I_3_’ and ‘I_4_’, respectively) formed between fully unfolded state (‘U’) and fully folded state (‘F’) in an apparently sequential order (Figure 2B and Supplementary Table S1). We first observed the hopping between U and I_1_ with an extension change of 4.4 ± 0.4 nm (mean ± S.E.) measured by Gaussian fitting. By decreasing the force to 11 pN, two other transitions (I_1_↔I_2_ and I_2_↔I_3_) with similar extension changes (4.9 ± 0.6 nm and 4.3 ± 0.5 nm, mean ± S.E.) appeared. Further decrease of the force to 9 pN gave rise to the transition I_3_↔I_4_ and I_3_→F with the extension changes of 3.9 ± 0.7 nm (mean ± S.E.) and 13.5 ± 0.2 nm (mean ± S.E.), respectively. Interestingly, once the tRNA folded into the ‘F’ state at 9 pN, no more extension changes were observed up to 30 seconds (data not shown).

The numbers of unfolded or folded nucleotides (Δ*N_ssRNA_*) during each transition were calculated according to the extension changes (see material and Methods). The Δ*N_ssRNA_* values of transitions U↔I_1_ and I_1_↔I_2_ were about 16-17 nt, corresponding to the nucleotide change of folding/unfolding a single stem-loop, and therefore we attributed the ‘I_1_’ and ‘I_2_’ states to the conformations with one and two stem-loops folded, respectively. The ‘I_3_’ state may represent a structure comprising three stem-loops, while the VL and the AS regions are still single-stranded under tension, which would thus leave 22 nucleotides unfolded. Alternatively, ‘I_3_’ state may represent a structure that two stem-loops are folded and coaxially stacked with each other (see Supplementary Text for more details). A transiently occupied state (‘I_4_’) with the extension 3.9 ± 0.7 nm shorter than ‘I_3_’ was typically seen, while no I_4_→F transition has been observed during all the force-clamp experiments. ‘I_4_’ probably represents a structure that the D stem, anti-codon stem and variable loop stack coaxially by formation of base triples C13-G22°G46, G10-C25°G45 and non-canonical base pair G26°A44, while U8°A14, U12-A23°A9, G15°C48 and the acceptor stem do not form (see Supplementary Text for more details).

### The folding/unfolding kinetics of TSL and DSL but not ASL depend on the existence of adjacent structures

Because the extension changes in forming each of the three stem-loops (i.e. ASL, TSL and DSL) are very close to each other, to further assign the transitions observed from force-clamp experiments, we extracted the force-dependent kinetics by applying Hidden Markov Chain analysis (42) to the extension trajectories of the wildtype tRNA^phe^ (Supplementary Table S2). We also performed the same measurements and extracted the kinetics as above for individual ASL, TSL and DSL molecules, respectively (Supplementary Table S3, S4). The results were plotted together and shown in Figure 2C. Interestingly, we found that the unfolding and folding kinetic characteristics of transition U↔I_1_ of the tRNA and the individual ASL are almost identical (Figure 2C). As a result, ‘I_1_’ can be unambiguously assigned to a structure in which only ASL was folded. However, the folding and unfolding kinetics of I 1↔I_2_ or I_2_↔I_3_ are different from those of individual TSL or DSL. The critical forces (i.e. the force at which the folded state and unfolded state are equally populated) of both I_1_↔I_2_ and I_2_↔I_3_ are larger than those of individual DSL and TSL (Figure 2C). Therefore, it is difficult to assign the I_1_↔I_2_ and I_2_↔I_3_ only according to the folding and unfolding kinetic information. ‘I_2_’ thus represented a structure in which either TSL or DSL could be folded besides the folded ASL. Based on the Δ*N_ssRNA_* values from the force-clamp experiments, the intermediate states in the single-molecule pulling experiments can also be assigned (Figure 2D, for detailed description, see Supplementary Text). The first (large) force rupture observed in the unfolding process was then found to correspond to the disruption of the major tertiary interactions of tRNA as well as other secondary structures (such as AS, TSL, DSL, etc.) except ASL all at once. The second unfolding rupture and the first refolding rupture were found to correspond to the unfolding/folding of ASL, in agreement with the U↔I_1_ transition in the force-clamp experiments.

In order to further understand how the inter-stem-loop interactions affect the folding and unfolding kinetics, we also measured the folding/unfolding kinetics of two truncated tRNA constructs DAV (contains DSL, ASL and VL) and AVT (contains ASL, VL and TSL) (Supplementary Figure S4, S5 and Supplementary Table S8-S11). Both constructs have only two apparent folding/unfolding transitions in either pulling or force-clamp experiments (Supplementary Figure S4). Remarkably, all the folding/unfolding kinetics of ASL in DAV and AVT are almost identical to those of U↔I_1_ in wildtype tRNA, further confirming that ‘I_1_’ corresponds to the structure that only ASL was folded. However, the folding and unfolding kinetics of DSL and TSL varies according to the sequence context. Interestingly, the critical forces, free energy differences and activation energies of DSL in DAV and TSL in AVT are 5.7 ± 0.7 and 2.5 ± 1.1 pN, and 3.3 ± 0.6 and 1.6 ± 0.4 kcal/mol, and 1.2 ± 0.5 and 0.7 ± 0.2 kcal/mol higher than those of individual DSL and TSL, respectively. These results suggested that some inter stem-loop interactions (for example, coaxial stacking) may stabilize the DAV and AVT structure, regardless of the entropy penalties introduced by junctional topological constraints (43). Moreover, the critical forces of DSL in DAV or TSL in AVT are also larger than those of I_1_↔I_2_ or I_2_↔I_3_ in the wildtype tRNA. These force-clamp experiment results demonstrated that the unfolding/folding kinetics of TSL and DSL in the tRNA are significantly affected by formation of other parts of the tRNA, probably due to the inter-stem-loop base stacking and steric constraints. The ASL formation, in sharp contrast, is always firstly folded and mostly isolated from the tertiary interactions, thus not affected by other parts of the tRNA.

### Mutant with disrupted tertiary interactions lost unfolding cooperativity and reduced unfolding energy barriers in the presence of Mg^2+^

To further investigate how tertiary interactions contribute to the cooperative unfolding of wildtype tRNA^phe^, we also measured the mechanical folding properties of a mutant G15/18/19A, in which the two base pairs in between D-loop and T-loop as well as the Levitt pair (G15°C48) were disrupted, as shown in Figure 3A. The typical pulling traces are shown in Figure 3B. In contrast to the unfolding pathway of the wildtype tRNA, the unfolding of the mutant yielded multiple intermediates (Figure 3B, blue lines) with the first unfolding transition occurring at lower forces. Moreover, the unfolding curve overlapped well with the refolding curves except for the first unfolding transition (Figure 3B, red lines), and the total nucleotides changed between the fully folded state and the fully unfolded state of the mutant is 73.2 ±1.6 nt (mean ± S.E., N = 12), which coincides with that of the wildtype tRNA. Such a finding suggests that disrupting the elbow structure leads to the loss of unfolding cooperativity. We denoted such an elbow-disrupted folded state as ‘F*’, to distinguish it from the canonical L-shape folded state ‘F’.

**Figure 3.**
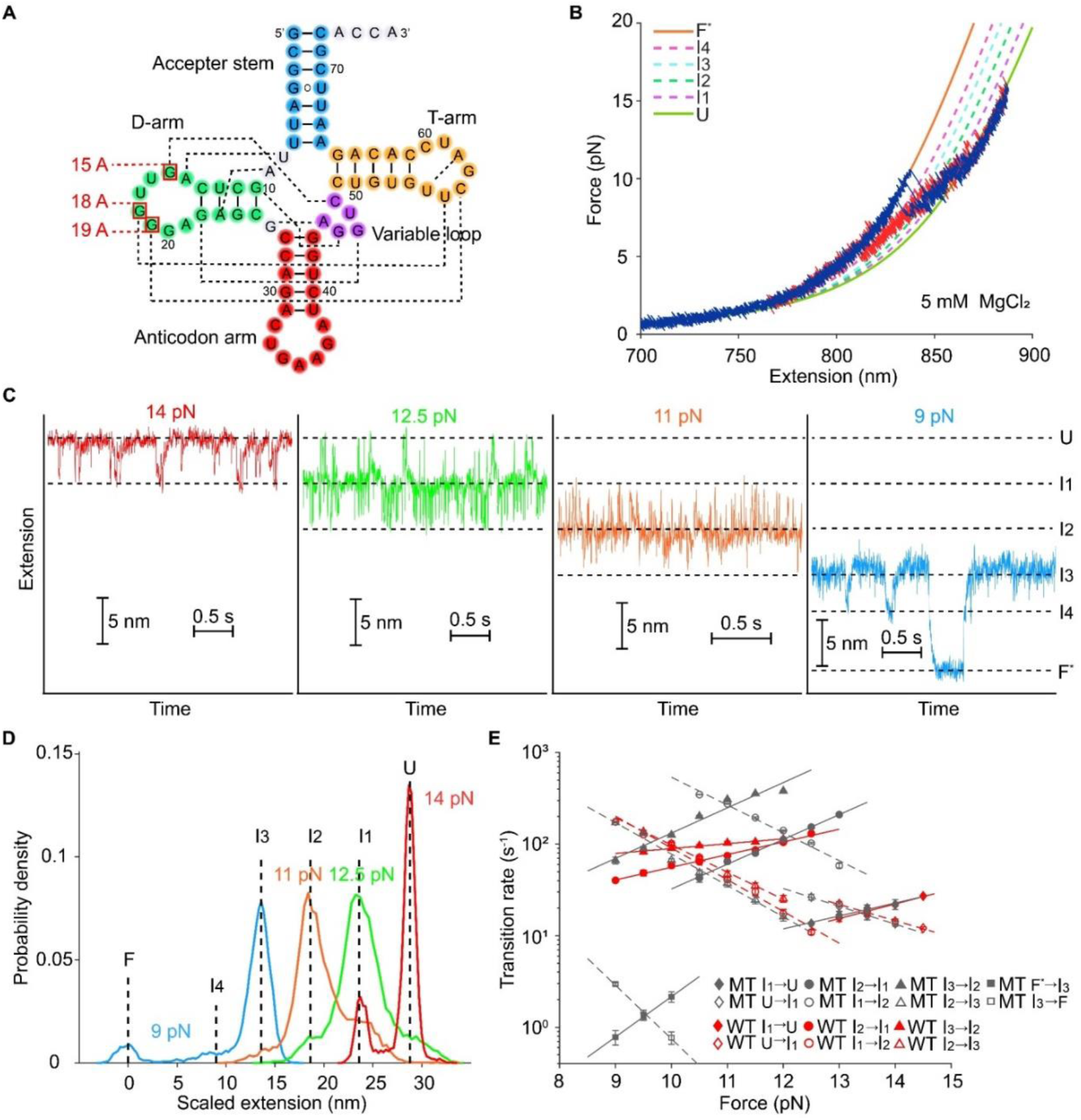
Mechanical unfolding and folding of mutant tRNA^phe^. (A) The secondary structure of a tRNA^phe^ three-nucleotide mutant where the guanine residues of position 15,18 and 19 are replaced with adenines, so that the two base pairs in between D-loop and T-loop as well as the Levitt pair are disrupted. (B) Typical FECs of unfolding (blue) and refolding (red) of the tRNA mutant during the pulling experiment at 5 mM MgCl_2_. (C) Typical force-dependent folding trajectories of the tRNA^phe^ mutant at 5 mM MgCl_2_. (D) Distribution of extension changes at different forces show force-dependent occupancies of the states. (E) Force-dependent kinetics of unfolding/folding tRNA^phe^ mutant at each transition during the force-clamp experiments.

In the force-clamp experiment, the unfolding cooperativity of mutant tRNA was also found to be significantly reduced compared to that of the wildtype (Figure 3C and Supplementary Figure S6B). Following the decrease of force from 14 pN to 11 pN, three transitions with similar extension changes occurred (Supplementary Table S5). Then, when the force dropped to 9 pN, the transiently occupied state ‘I_4_’ could also be observed. It is interesting that besides the reversible transition of I_3_↔I_4_ at 9 pN, another reversible transition with a large extension changed (13.2 ± 0.2 nm, mean ± S.E., coincided with the I_3_→F transition of the wildtype at 9 pN) was also present, indicating that it is a transition between the ‘I_3_’ and ‘F*’. These results suggest that the disruption of the tertiary interactions of the elbow region destabilizes the folded tRNA and thus encourages the transition between the ‘I_3_’ and ‘F*’.

To further unravel how the disruption of the tertiary interactions in the elbow region affect the folding and unfolding behaviors of the stem-loops, we also extracted the force-dependent kinetics using the Hidden Markov Chain analysis (42) (Figure 3E and Supplementary Table S6). Obviously, the disruption of the tertiary interactions in the elbow region significantly affected the folding and unfolding kinetics of I_2_↔I_3_ and I_1_↔I_2_ but not U↔I_1_. These results demonstrate that besides the junctional topological constraints (44) (as the G15/18/19A mutation does not disrupt the junctional bases of the tRNA), the tertiary interactions in the elbow region also play an important role in tuning the folding kinetics of the tRNA.

To better illustrate the critical roles of the tertiary interactions in the elbow region, we also reconstructed the free energy landscapes of the folding and unfolding pathway for both wildtype and mutant tRNA. We applied the Jarzynski’s equality to the FECs to determine the free energy change of the F→I_1_ transition of the wildtype tRNA (45,46), and the position of the transition state along the reaction coordinate and transition barrier height were extracted by fitting the rupture force distributions to the Dudko-Hummer-Szabo model (see material and methods) (24). These results were combined with the force-clamp results for all the other transitions, to reconstruct the piecewise energy landscape for the unfolding and folding pathway of both the wildtype and mutant tRNA^phe^ at 10 pN, with 5 mM MgCl_2_ (Figure 4). Comparing the folding/unfolding energy landscape of wildtype and mutant tRNA^phe^, we found that the free energy of ‘I_1_’ and the barrier of I_1_→U of mutant tRNA^phe^ align quite well with its wildtype, while the free energy of ‘F*’ state for the mutant is about 5.8 ± 2.0 kcal/mol higher than the ‘F’ state for the wildtype tRNA at 10 pN. Remarkably, the barriers of I_2_→I_1_ and I_3_→I_2_ of mutant tRNA^phe^ are lower and locate closer to the unfolded states when the elbow structures are disrupted. The free energies of ‘I_2_’ and ‘I_3_’ also slightly decreased with the disruption of the elbow structure. As I_2_→I_1_ and I_3_→I_2_ correspond to the unfolding of DSL and/or TSL, these results suggest that tertiary interactions in the elbow region destabilize the secondary structures of D stem and/or T stem probably by driving their conformations away from canonical A-form helix, and/or by reducing the flexibility of the whole molecule (47). On the other hand, these tertiary interactions increased the free energies of not only the largest cooperative transition barrier but also the barriers between intermediate states, suggesting the key roles of tertiary interaction network in tRNA unfolding. These results clearly demonstrated that the mechanical unfolding of secondary structure and tertiary structure of the tRNA are coupled.

**Figure 4.**
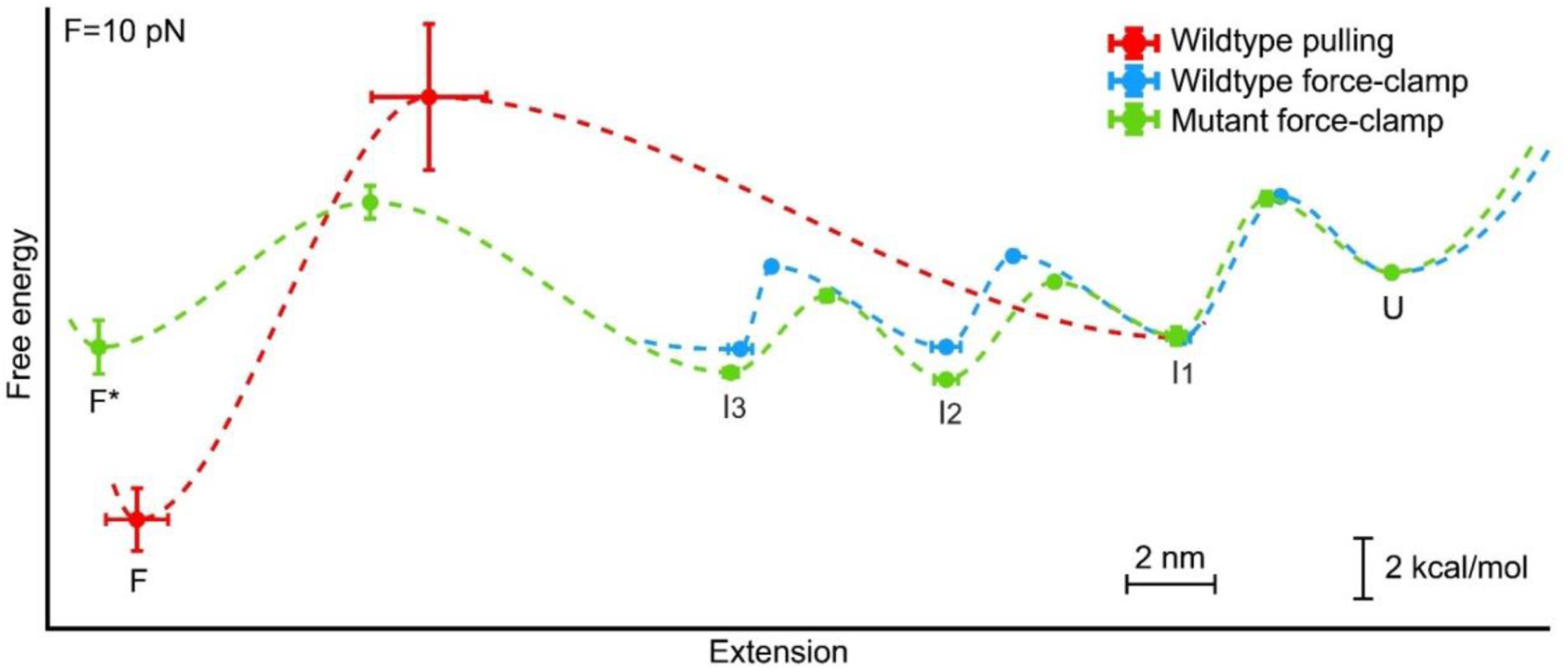
Free energy landscapes for folding/unfolding of the wildtype and mutant tRNA^phe^ at 10 pN at 5 mM MgCl_2_. The key features of the energy landscapes for the five-state unfolding pathway were reconstructed from piecewise two-state analyses of each transition. Energies and positions are plotted with reference to the ‘U’ state. Error bars are presented as S.E.M. Dotted lines indicate notional landscape shapes in the presence of 5 mM MgCl_2_ for the wildtype (red for pulling experiment and blue for force-clamp experiment) and mutant (green).

### Unfolding of wildtype tRNA^phe^ involves multiple steps and is largely reversible with increased unfolding rates in low-concentration Mg^2+^ solution

As tertiary interactions such as G18-U55 and G19-C56 base pairings depend on Mg^2+^ concentrations (14,16,48,49), we then investigated how Mg^2+^ concentration affects the folding/unfolding of the wildtype tRNA via pulling experiments. The typical FECs measured at 5 mM, 2 mM and 0.5 mM MgCl_2_ are shown in Figure 5A-C, respectively. For all three concentrations, the refolding traces of FECs showed a multi-step feature similar to those described above (data not shown). At 5 mM and 2 mM MgCl_2_ conditions (Figure 5A, B), all the unfolding FECs exhibited the typical two-step features similar to Fig. 2D, indicating that the tRNA was initially in the fully folded canonical L shape when no external force was exerted. However, at 0.5 mM MgCl_2_, we also observed that about 8% (5/65) of unfolding events had multiple intermediates (Figure 5C, blue line) with the first unfolding transition occurred at lower forces, which is similar to the unfolding pathway of the mutant tRNA at 5 mM MgCl_2_ (Figure 3B). Such a finding suggests that at 0.5 mM MgCl_2_, the elbow region of a small portion of wildtype tRNA molecule may be disrupted before the mechanical pulling process, namely an equilibrium between states ‘F’ and ‘F*’ may exist at 0.5 mM MgCl_2_ when no external force was exerted, which is consistent with previous studies (15,16). To exclude the possibility that such a small fraction of traces may come from tether heterogeneity, we also tried to measure the same tether multiple times. As can be seen in Figure 5D, even for the same tether, at 0.5 mM MgCl_2_ conditions, sometimes multi-step unfolding can be seen, which hence rules out that such a phenomenon is due to tether-tether variation. Moreover, the unfolding curve overlapped well with the refolding curves except for the first unfolding transition, further suggesting that these small fractions of wildtype tRNA molecules may fold into the ‘F*’ state. The force-dependent kinetic rates of F→I_1_ at different concentrations of MgCl_2_ were also calculated using the Dudko-Hummer-Szabo method (see material and methods) (24), as shown in Figure 5E and Supplementary Table S7. It showed that the transition rate of F→I_1_ at the same force was enhanced by almost one order when Mg^2+^ concentration decreased from 5 mM to 0.5 mM. All these results highlight the significant role of Mg^2+^ in decreasing tRNA unfolding rates and stabilizing its canonical structure.

**Figure 5.**
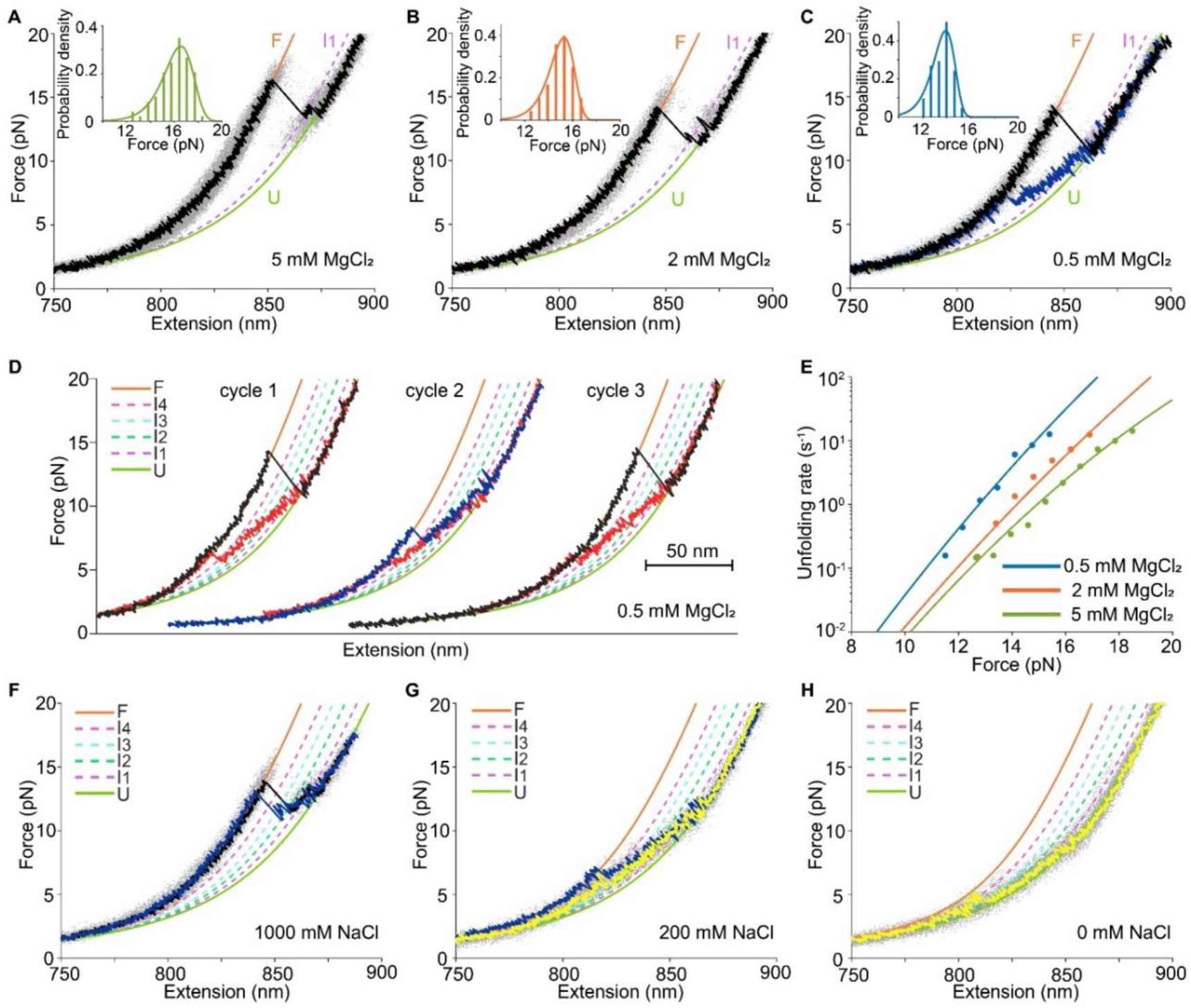
Mechanical unfolding and folding of wildtype tRNA^phe^ at different salt conditions. (A-C) FECs at 5 mM, 2 mM and 0.5 mM MgCl_2_, respectively. Typical FECs are plotted above the aggregated data from over 50 FECs (grey dots). Black curves represent typical unfolding trajectories starting from canonical L-shape tRNA (‘F’) and blue curves represent typical unfolding trajectories starting from a structure in which all the stem-loops were folded but the base pairs between D-loop and T-loop were disrupted (‘F*’). The insets showed the distributions of rupture forces of the F→I_1_ transition. (D) FECs of three pulling cycles of a single tether at 0.5 mM MgCl_2_. Black curves represent the unfolding half-cycle starting from the ’F’ state and the blue curve represents an unfolding half-cycle starting from the ‘F*’ state. Red curves represent the refolding half-cycles. (E) Force-dependent unfolding rates of F→I_1_ transition at 5 mM, 2 mM and 0.5 mM MgCl_2_, respectively. (F-H) FECs of unfolding trajectories at 1000 mM, 200 mM and 0 mM NaCl, respectively. Typical FECs are plotted above the aggregated data. Black, blue and yellow curves represent the unfolding trajectories starting from ‘F’, ‘F*’ and probable ‘I_4_’ state, respectively.

### Unfolding of wildtype tRNA^phe^ initiates from different structures in different concentrations of NaCl solution

It has been controversial whether the unmodified yeast tRNA^phe^ can fold into the canonical L-shape structure in buffers containing Na^+^ instead of Mg^2+^ (16,48–50). To clarify whether the unmodified yeast tRNA^phe^ still unfolds cooperatively in buffers containing only Na^+^, we measured the unfolding of single wildtype tRNA^phe^ molecule at different NaCl concentrations in the absence of Mg^2+^. As shown in Figure 5F, at 1000 mM NaCl, 34% (85/253) of unfolding events are non-cooperative with multiple intermediates, while the other 66% unfold cooperatively as most unfolding events at MgCl_2_, suggesting that a large portion of the unmodified wildtype tRNA is still capable of folding into the L-shape structure at 1000 mM NaCl when no external force was exerted.

At 200 mM NaCl, 51% (34/67) of traces unfolded non-cooperatively with multiple intermediates and overlapped well with the refolding curves, similar to the small fraction of traces at 0.5 mM MgCl_2_ (Figure 5G, blue line). Interestingly, we found that the other 49% of traces started unfolding neither from the state ‘F’ nor ‘F*’ but from a state whose FECs can be well aligned with the intermediate ‘I_4_’ (Figure 5G, yellow line), suggesting that the acceptor stem may not form in low-salt buffers even when no external force was exerted. Such an observation agrees with previous temperature and urea denaturation experiments (16,19). Once NaCl concentration went down to zero, almost all of the traces (66/69) started unfolding from a state in which AS was probably not folded (see Discussion). The non-cooperative unfolding and refolding of these traces also suggest that the elbow structure did not form in this state.

### Molecular dynamics simulation results further suggest tertiary interactions in the elbow region dominate the overall cooperativity and increase energy barriers of the tRNA unfolding pathway

To further illustrate the mechanism of tRNA unfolding at the atomic level, we applied steered molecular dynamics (SMD) simulations based on Martini coarse-grained (CG) model, to visualize the entire structural dynamics of the tRNA unfolding in details (Supplementary Figure S7 and Movie S1). We compared the CG SMD simulations initiating from a canonical L-shaped folded tRNA (‘F’, Fig. 6A) with those initiating from an elbow-disrupted folded tRNA (‘F*’, Figure 6B) to investigate the unfolding behaviors including the structural transformation, the force-extension response, and the extension spans indicating the onset and the end of stem-loop disruptions (Figure 6C, D). For the canonical L-shape folded state (‘F’) in the presence of Mg^2+^, the secondary structures were unfolded sequentially in the order of AS→TSL→DSL→ASL (Figure 6C and Supplementary Figure S7). This is consistent with the stepwise unfolding trajectories of force-clamp experiments, in which the AS unfolds first and the ASL unfolds last. We found that the tRNA unfolding is kept sequential and the unfolding sequence is the same when the pulling rates changes from 0.2 to 10 nm/ns (Supplementary Figure S8), suggesting that the pulling rate does not affect the unfolding sequence of the secondary structures of the tRNA. Although the secondary structures of the tRNA unfold separately during the SMD, the unfolding of secondary and tertiary structures are still coupled (Supplementary Movie S1). To overcome the energy barrier for starting up the disruption of each secondary structure, the force-extension profile showed a high level of unfolding force with multiple peaks. The similar unfolding sequence and profile were observed in the case without Mg^2+^ (Supplementary Figure S9), which suggests that the canonical L-shape tRNA undergoes the sequential unfolding regardless of the Mg^2+^ concentrations. However, the unfolding of tRNA beginning with the elbow-disrupted folded state (‘F*’) showed a different scenario in the force-extension profile and the structural disruptions (Figure 6D and Supplementary Movie S2). The profile exhibited a plateau without pronounced peaks during the disruptions of TSL, DSL and ASL. Meanwhile, the overlapped extension spans indicated the disruptions of the secondary structures are partially coupled. Without the initiation barrier, the successive disruptions can lower the level of the unfolding force and induce the continuous unfolding, which is consistent with the pulling experiment results of the mutant tRNA in the presence of Mg^2+^ and those of the wildtype tRNA in solutions with low-concentration or no Mg^2+^. These results led to a conclusion that the unfolding behaviors of tRNA directly depended on the starting folding states and were highly correlated to the tertiary structures. To study the effects of ions, we adopted all-atom simulations and found that more energies are required for the disruptions of the elbow and the AS structures with Mg^2+^ or Na^+^ ions suggesting the cations can facilitate the formation and stability of the canonical L-shape folded structure (‘F’) (Supplementary Figure S10, S11). These findings are consistent with our pulling and force-clamp experiment results as well that disrupting the elbow structure significantly lowers the unfolding barriers.

**Figure 6.**
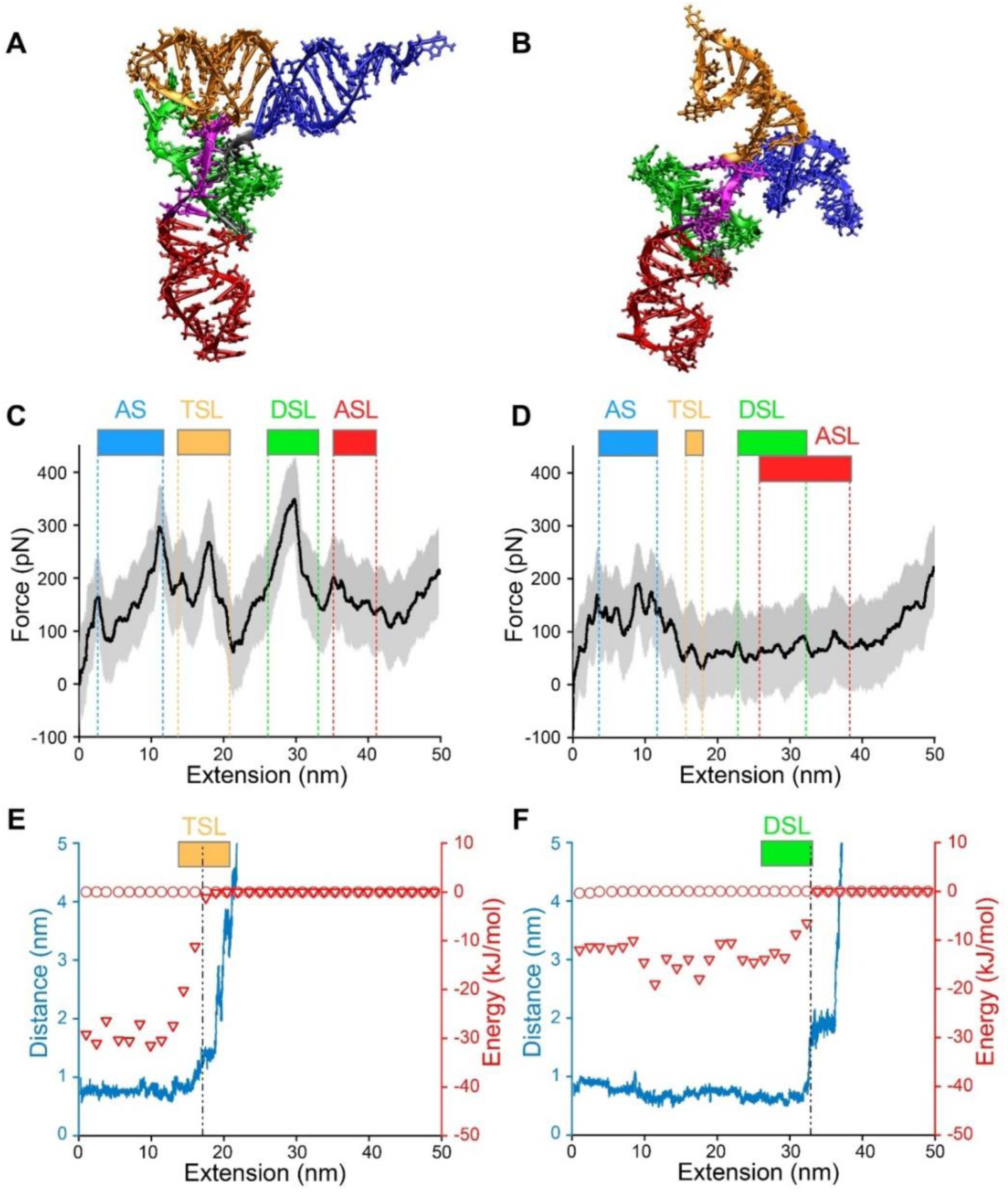
SMD simulations reveal the cation effect and the unfolding cooperativity. (A, B) Starting SMD configurations of the canonical L-shape folded state ‘F’ (A) and the elbow-disrupted folded state ‘F*’ (B). (C, D) FECs by SMD simulations initiating from the ‘F’ state with Mg^2+^ (C) and the ‘F*’ state without Mg^2+^ (D) at the pulling rate of 0.2 nm/ns. The black lines represent the running averages with a window size of 400 ps. The gray shades denote the standard deviations of the running averaging. Colored bars and dashed lines represent the extension spans indicating the onset and the end of stem-loop disruptions (AS, blue; TSL, yellow; DSL, green; ASL, red). (E, F) The distances and binding energies of the G19-C56 (E) and G22-C46 (F) base pairs as a function of extension. The blue lines represent the distance-extension curves of the starting ‘F’ structure. The red triangles show binding energies for the starting ‘F’ structure, while the red circles for the starting ‘F*’ structure. The dash-dot lines mark the critical breaking of the base pairs.

To further study the coupling between the secondary and tertiary structures during the tRNA unfolding, we reviewed the disruptions of elbow base pairs (for example, G19-C56) and D stem base pairs (for example, G22-C46) in the CG SMD simulations. The distances and the binding energies of the two base pairs were calculated for the two starting folding states as shown in Figure 6E, F. Unlike the base pairs that remained no interactions (zero binding energies) in the case of the elbow-disrupted folded structure, G19-C56 and G22-G46 became unpaired during the unfolding of the canonical L-shape tRNA indicated by the surging of the distance and the energy. Moreover, the dissociations of the G19-C56 and G22-C46 base pairs were strongly coupled with the disruptions of the TSL and the DSL, respectively. Combining the SMD simulations with experiment results, we have revealed that formation of the elbow region plays a critical role in determining the overall unfolding stability and cooperativity of tRNA.

## DISCUSSION

In this work, we unraveled the mechanical unfolding and folding details of unmodified yeast cytosolic tRNA^phe^. Our results clearly showed that the mechanical unfolding and folding of secondary and tertiary structures of tRNA are coupled due to tertiary interactions. Based on the experiment and simulation results as well as some assumptions for ‘I_3_’ and ‘I_4_’ (see Supplementary Text), we summarized the folding and unfolding pathways of tRNA^phe^ as follows (Figure 7).

**Figure 7.**
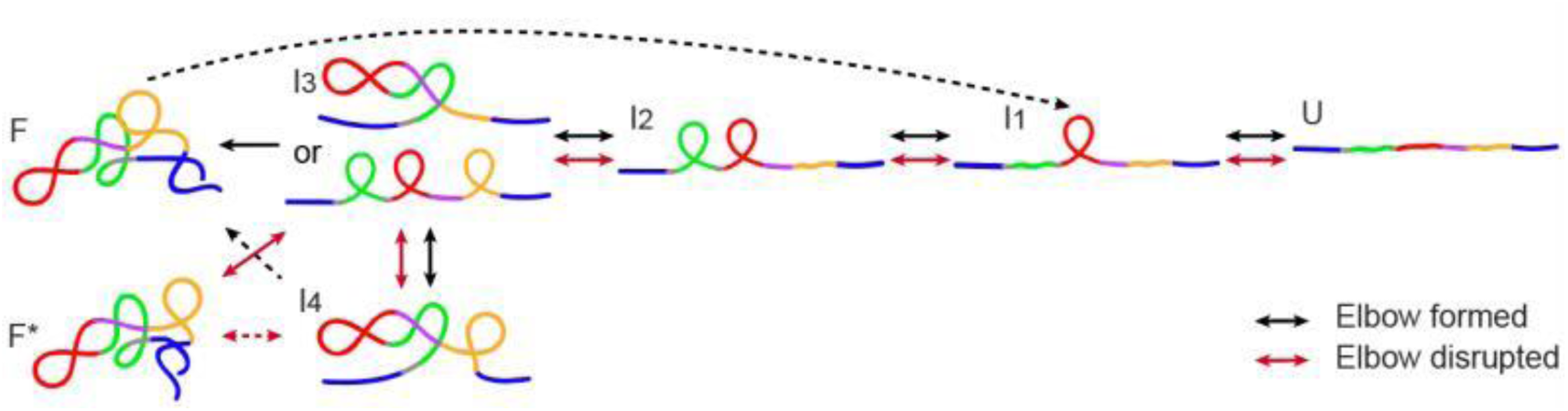
Summary of the folding and unfolding pathways of tRNA^phe^. When the elbow is formed (wildtype at 5 mM MgCl_2_ or 1000 mM NaCl), the L-shape structure tRNA^phe^ (‘F’) unfolds in two steps and refolds in multiple steps (black arrows). When the elbow is disrupted (wildtype at 0.5 mM MgCl_2_, 200mM NaCl or mutant at 5 mM MgCl_2_) (‘F*’), the tRNA unfolds and refolds less cooperatively in multiple steps (red arrows). Solid arrows indicate transitions observed in force-clamp experiments, and dashed arrows indicate transitions observed in pulling experiments. More detailed pathways can be found in SI Supplementary Figure S12.

For the biological implications, our studies suggest that the folding and unfolding of tertiary structures and the secondary structures of the tRNA are also coupled at physiological conditions. The widely proposed cloverleaf-shape intermediate structure may not exist in the tRNA folding or unfolding pathway at physiological conditions even when no external force is exerted, as inter-stem-loop interactions such as coaxial stacking and long-range base pairing may exist even when the AS are not folded. Interestingly, such cooperativity also exists in the mechanical unfolding of a tRNA mimic structure termed TSS within the 3’ UTR of Turnip crinkle virus genome (51). Magnesium ions stabilize the tRNA mimic structure and increase its overall unfolding and folding cooperativity, although its secondary structure and tertiary interactions are completely different from a canonical tRNA (51). Intriguingly, the stabilization effect of magnesium ions was found to be asymmetric for tRNA folding and unfolding processes in our experiments. The presence of magnesium ions did not significantly increase the mechanical folding cooperativity of yeast tRNA^phe^ in our pulling experiments, although it is known that they increase the thermal folding/unfolding cooperativity in the bulk studies. This result thus highlights the difference between the mechanical and thermal folding/unfolding processes.

Our studies may explain how the two tRNA chaperones *E. coli* TruB and TrmA solved the kinetically trapped partially folded and misfolded tRNA in the absence of ATP hydrolysis activity. Tertiary interactions participating in forming the elbow region maintain the coupling between secondary and tertiary structures and increase most energy barriers during the unfolding pathway. Although the partially folded or misfolded tRNA may not have a complete elbow structure, such coupling effect may still exist once a portion of these tertiary interactions remain. Once disrupting these interactions, tRNA-chaperone binding may decouple the unfolding of secondary and tertiary structures of the partially folded and misfolded tRNA, resulting in significantly lowering the unfolding cooperativity as well as the energy barriers, and thus increasing unfolding rates of the partially folded and misfolded tRNA. Our findings suggest that tRNA chaperone activity maybe a common feature of tRNA binding proteins which disrupt the tertiary interactions of tRNA in the elbow region.

Although the mechanical unfolding of the tRNA is highly cooperative, the folding and unfolding of ASL are independent of other parts of the tRNA. Such a mechanical property is consistent with the fact that some modifications on ASL do not require the existence of the elbow structure (52). We suggest the nascent pre-tRNA may have similar mechanical folding and unfolding pathways to the matured tRNA, namely the folding and unfolding of the pre-tRNA secondary and tertiary structures are highly cooperative except for the anti-codon-intron stem-loop.

In summary, we unraveled the detailed mechanical folding and unfolding mechanisms of the unmodified yeast tRNA^phe^ under near physiological conditions, which shall serve as a cornerstone for studying mechanical properties of other types of tRNAs as well as tRNAs with nucleotide modifications. These results shall help gain more insights into how the mechanical stability of tRNA affects its interactions with proteins and complexes and hence its biological activities.

## AVAILABILITY

All data that support the findings of this study are available from the corresponding authors on request.

## SUPPLEMENTARY DATA

Supplementary Data are available at NAR online.

## ACKNOWLEDGEMENT

We thank Associate Professor Gang Chen at CUHK-Shenzhen for insightful discussion and suggestions. We thank Assistant Professor Bo Sun at ShanghaiTech University for providing 0.5-μm anti-digoxigenin coated polystyrene beads. Part of the experiments reported here was conducted on the Physical Research Platform in School of Physics, Sun Yat-sen University (PRPSP, SYSU) and the simulations reported were performed on resources provided by the National Supercomputer Center in Guangzhou.

## REFERENCES

1. Hasler, D. and Meister, G. (2016) From tRNA to miRNA: RNA-folding contributes to correct entry into noncoding RNA pathways. FEBS Lett., 590, 2354–2363.

2. Hasler, D., Lehmann, G., Murakawa, Y., Klironomos, F., Jakob, L., Grässer, Friedrich A., Rajewsky, N., Landthaler, M. and Meister, G. (2016) The lupus autoantigen la prevents mis-channeling of tRNA fragments into the human microRNA pathway. Mol. Cell, 63, 110–124.

3. Whipple, J.M., Lane, E.A., Chernyakov, I., D’Silva, S. and Phizicky, E.M. (2011) The yeast rapid tRNA decay pathway primarily monitors the structural integrity of the acceptor and T-stems of mature tRNA. Genes. Dev., 25, 1173–1184.

4. Lan, P., Tan, M., Zhang, Y., Niu, S., Chen, J., Shi, S., Qiu, S., Wang, X., Peng, X., Cai, G. et al. (2018) Structural insight into precursor tRNA processing by yeast ribonuclease P. Science, 362, eaat6678.

5. Megel, C., Morelle, G., Lalande, S., Duchêne, A.-M., Small, I. and Maréchal-Drouard, L. (2015) Surveillance and cleavage of eukaryotic tRNAs. Int. J. Mol. Sc.i, 16, 1873–1893.

6. Flanagan, J.F., Namy, O., Brierley, I. and Gilbert, R.J.C. (2010) Direct observation of distinct A/P hybrid-state tRNAs in translocating ribosomes. Structure, 18, 257–264.

7. Wang, X., Jia, H., Jankowsky, E. and Anderson, J.T. (2008) Degradation of hypomodified tRNA(iMet) in vivo involves RNA-dependent ATPase activity of the DExH helicase Mtr4p. RNA, 14, 107–116.

8. Chimnaronk, S., Suzuki, T., Manita, T., Ikeuchi, Y., Yao, M., Suzuki, T. and Tanaka, I. (2009) RNA helicase module in an acetyltransferase that modifies a specific tRNA anticodon. EMBO J., 28, 1362–1373.

9. Keffer-Wilkes, L.C., Veerareddygari, G.R. and Kothe, U. (2016) RNA modification enzyme TruB is a tRNA chaperone. Proc. Natl. Acad. Sci. U. S. A., 113, 14306–14311.

10. Keffer-Wilkes, L.C., Soon, E.F. and Kothe, U. (2020) The methyltransferase TrmA facilitates tRNA folding through interaction with its RNA-binding domain. Nucleic Acids Res., 48, 7981–7990.

11. Porat, J., Kothe, U. and Bayfield, M.A. (2021) Revisiting tRNA chaperones: New players in an ancient game. RNA, 27, 543–559.

12. Serebrov, V., Clarke, R.J., Gross, H.J. and Kisselev, L. (2001) Mg2+-induced tRNA folding. Biochemistry, 40, 6688–6698.

13. Privalov, P.L. and Filimonov, V.V. (1978) Thermodynamic analysis of transfer RNA unfolding. J. Mol. Biol., 122, 447–464.

14. Behlen, L.S., Sampson, J.R., DiRenzo, A.B. and Uhlenbeck, O.C. (1990) Lead-catalyzed cleavage of yeast tRNAPhe mutants. Biochemistry, 29, 2515–2523.

15. Shelton, V.M., Sosnick, T.R. and Pan, T. (1999) Applicability of urea in the thermodynamic analysis of secondary and tertiary RNA folding. Biochemistry, 38, 16831–16839.

16. Shelton, V.M., Sosnick, T.R. and Pan, T. (2001) Altering the intermediate in the equilibrium folding of unmodified yeast tRNAPhe with monovalent and divalent cations. Biochemistry, 40, 3629–3638.

17. Beltchev, B., Yaneva, M. and Staynov, D. (1976) Thermal melting curves of tRNAPhe from yeast lacking different numbers of nucleotides from the 3’-end. Eur. J. Biochem., 64, 507–510.

18. Mustoe, A.M., Brooks, C.L., III and Al-Hashimi, H.M. (2014) Topological constraints are major determinants of tRNA tertiary structure and dynamics and provide basis for tertiary folding cooperativity. Nucleic Acids Res., 42, 11792–11804.

19. Leamy, K.A., Yamagami, R., Yennawar, N.H. and Bevilacqua, P.C. (2019) Single-nucleotide control of tRNA folding cooperativity under near-cellular conditions. Proc. Natl. Acad. Sci. U. S. A., 116, 23075–23082.

20. Voigts-Hoffmann, F., Hengesbach, M., Kobitski, A.Y., van Aerschot, A., Herdewijn, P., Nienhaus, G.U. and Helm, M. (2007) A methyl group controls conformational equilibrium in human mitochondrial tRNALys. J. Am. Chem. Soc., 129, 13382–13383.

21. Kobitski, Andrei Y., Hengesbach, M., Seidu-Larry, S., Dammertz, K., Chow, Christine S., van Aerschot, A., Nienhaus, G.U. and Helm, M. (2011) Single-molecule FRET reveals a cooperative effect of two methyl group modifications in the folding of human mitochondrial tRNALys. Chem. Biol., 18, 928–936.

22. Smith, A.M., Abu-Shumays, R., Akeson, M. and Bernick, D.L. (2015) Capture, Unfolding, and Detection of Individual tRNA Molecules Using a Nanopore Device. Front. Bioeng. Biotechnol., 3, 91.

23. Palassini, M. and Ritort, F. (2011) Improving free-energy estimates from unidirectional work measurements: Theory and experiment. Phys. Rev. Lett., 107, 060601.

24. Dudko, O.K., Hummer, G. and Szabo, A. (2006) Intrinsic rates and activation free energies from single-molecule pulling experiments. Phys. Rev. Lett., 96, 108101.

25. Zhu, W., von dem Bussche, A., Yi, X., Qiu, Y., Wang, Z., Weston, P., Hurt, R.H., Kane, A.B. and Gao, H. (2016) Nanomechanical mechanism for lipid bilayer damage induced by carbon nanotubes confined in intracellular vesicles. Proc. Natl. Acad. Sci. U. S. A., 113, 12374–12379.

26. Anthony, P.C., Perez, C.F., García-García, C. and Block, S.M. (2012) Folding energy landscape of the thiamine pyrophosphate riboswitch aptamer. Proc. Natl. Acad. Sci. U. S. A., 109, 1485–1489.

27. Gál, J. and Kálmán, M. (2002) Autosticky PCR. Directional cloning of PCR products with performed 5’ overhangs. Methods Mol. Biol., 192, 141–151.

28. Liu, X.-P. and Liu, J.-H. (2010) The terminal 5’phosphate and proximate phosphorothioate promote ligation-independent cloning. Protein Sci., 19, 967–973.

29. Sayers, J.R., Schmidt, W. and Eckstein, F. (1988) 5’ - 3’exonucleases in phosphorothioate-based oligonucleotide-directed mutagenesis. Nucleic Acids Res., 16, 791–802.

30. Yang, D., Liu, W., Deng, X., Xie, W., Chen, H., Zhong, Z. and Ma, J. (2020) GC-Content Dependence of Elastic and Overstretching Properties of DNA:RNA Hybrid Duplexes. Biophys. J., 119, 852–861.

31. Wang, M.D., Yin, H., Landick, R., Gelles, J. and Block, S.M. (1997) Stretching DNA with optical tweezers. Biophys. J., 72, 1335–1346.

32. Abraham, M.J., Murtola, T., Schulz, R., Páll, S., Smith, J.C., Hess, B. and Lindahl, E. (2015) GROMACS: High performance molecular simulations through multi-level parallelism from laptops to supercomputers. SoftwareX, 1-2, 19–25.

33. Maier, J.A., Martinez, C., Kasavajhala, K., Wickstrom, L., Hauser, K.E. and Simmerling, C. (2015) ff14SB: Improving the Accuracy of Protein Side Chain and Backbone Parameters from ff99SB. J. Chem. Theory Comput., 11, 3696–3713.

34. Pérez, A., Marchán, I., Svozil, D., Sponer, J., Cheatham, T.E., Laughton, C.A. and Orozco, M. (2007) Refinement of the AMBER Force Field for Nucleic Acids: Improving the Description of α/γ Conformers. Biophys. J., 92, 3817–3829.

35. Zgarbová, M., Otyepka, M., Šponer, J., Mládek, A., Banáš, P., Cheatham, T.E. and Jurečka, P. (2011) Refinement of the Cornell et al. Nucleic Acids Force Field Based on Reference Quantum Chemical Calculations of Glycosidic Torsion Profiles. J. Chem. Theory Comput., 7, 2886–2902.

36. Jorgensen, W.L., Chandrasekhar, J., Madura, J.D., Impey, R.W. and Klein, M.L. (1983) Comparison of simple potential functions for simulating liquid water. J. Chem. Phys., 79, 926–935.

37. Essmann, U., Perera, L., Berkowitz, M.L., Darden, T., Lee, H. and Pedersen, L.G. (1995) A smooth particle mesh Ewald method. J. Chem. Phys., 103, 8577–8593.

38. Isralewitz, B., Gao, M. and Schulten, K. (2001) Steered molecular dynamics and mechanical functions of proteins. Curr. Opin. Struct., 11, 224–230.

39. Kim, W., Zhu, W., Hendricks, G.L., Van Tyne, D., Steele, A.D., Keohane, C.E., Fricke, N., Conery, A.L., Shen, S., Pan, W. et al. (2018) A new class of synthetic retinoid antibiotics effective against bacterial persisters. Nature, 556, 103–107.

40. Uusitalo, J.J., Ingólfsson, H.I., Marrink, S.J. and Faustino, I. (2017) Martini Coarse-Grained Force Field: Extension to RNA. Biophys. J., 113, 246–256.

41. Tironi, I.G., Sperb, R., Smith, P.E. and Gunsteren, W.F.v. (1995) A generalized reaction field method for molecular dynamics simulations. J. Chem. Phys., 102, 5451–5459.

42. van de Meent, J.-W., Bronson, Jonathan E., Wiggins, Chris H. and Gonzalez, Ruben L. (2014) Empirical bayes methods enable advanced population-level analyses of single-molecule FRET experiments. Biophys. J., 106, 1327–1337.

43. Mak, C.H. and Phan, E.N.H. (2018) Topological Constraints and Their Conformational Entropic Penalties on RNA Folds. Biophys. J., 114, 2059–2071.

44. Crothers, D.M., Cole, P.E., Hilbers, C.W. and Shulman, R.G. (1974) The molecular mechanism of thermal unfolding of Escherichia coli formylmethionine transfer RNA. J. Mol. Biol., 87, 63–88.

45. Liphardt, J., Dumont, S., Smith, S.B., Tinoco, I. and Bustamante, C. (2002) Equilibrium information from nonequilibrium measurements in an experimental test of Jarzynski’s equality. Science, 296, 1832–1835.

46. Gore, J., Ritort, F. and Bustamante, C. (2003) Bias and error in estimates of equilibrium free-energy differences from nonequilibrium measurements. Proc. Natl. Acad. Sci. U. S. A., 100, 12564–12569.

47. Friederich, M.W., Vacano, E. and Hagerman, P.J. (1998) Global flexibility of tertiary structure in RNA: Yeast tRNAPhe as a model system. Proc. Natl. Acad. Sci. U. S. A., 95, 3572–3577.

48. Maglott, E.J., Deo, S.S., Przykorska, A. and Glick, G.D. (1998) Conformational transitions of an unmodified tRNA: implications for RNA folding. Biochemistry, 37, 16349–16359.

49. Hall, K.B., Sampson, J.R., Uhlenbeck, O.C. and Redfield, A.G. (1989) Structure of an unmodified tRNA molecule. Biochemistry, 28, 5794–5801.

50. Friederich, M.W. and Hagerman, P.J. (1997) The angle between the anticodon and aminoacyl acceptor stems of yeast tRNAPhe is strongly modulated by magnesium ions. Biochemistry, 36, 6090–6099.

51. Camunas-Soler, J., Alemany, A. and Ritort, F. (2017) Experimental measurement of binding energy, selectivity, and allostery using fluctuation theorems. Science, 355, 412–415.

52. Jiang, H.-Q., Motorin, Y., Jin, Y.-X. and Grosjean, H. (1997) Pleiotropic effects of intron removal on base modification pattern of yeast tRNAPhe: an in vitro study. Nucleic Acids Res., 25, 2694–2701.

