## Supplemental Information for "Single-molecule mechanical unfolding kinetics of unmodified *Saccharomyces cerevisiae* tRNA^Phe^: a hint to the tRNA chaperone-tRNA interaction mechanism"

**This PDF file includes:**

Supplementary text
Figures S1 to S12
Legends for Movies S1 to S2
Tables S1 to S12
References

**Other supplementary materials for this manuscript include the following:**

Movies S1 to S2

### Supplementary Text

#### The temporal resolution of dual-trap optical tweezers

The measurable kinetics of force-clamp experiments are limited by the autocorrelation time ($\tau_0$), which is defined as $\tau_0$ =1/2 π$f_c$ (1). The autocorrelation function (ACF) for the time series of force signal, $F_i$, with sampling rate $\delta$ and $N$ data points is calculated as,

$$CF\left(\tau = \frac{j}{\delta}\right) = \frac{\sum_{i=0}^{N-1}(F_i - \mu)\left(F_{1-j} - \mu\right)}{\sum_{i=0}^{N-1}(F_i - \mu)^2} \tag{1}$$

where $\mu$ is the mean of $F$ signal. A typical ACF was shown in Supplementary Figure S2. The plot is fitted by a single exponential decay function,

$$ACF(\tau) = \exp\left(\frac{\tau}{\tau_0}\right) \tag{2}$$

where $\tau_0$ is the fitted parameter. The average $\tau_0$ from 10 beads was 0.041 ± 0.002 ms (mean ± S.E.), which implies that it was possible to measure accurately the RNA folding kinetics as fast as 0.041 ms in our force-clamp experiments using dual-trap optical tweezers. All the measured kinetics in our force-clamp experiments are at least two orders of magnitude slower than $\tau_0$.

#### Data analysis

*Modified WLC fitting*

The extension trajectories obtained from force-clamp experiments were smoothed to 20 kHz, and then the extension change for each transition, $\Delta x$, was measured directly from Gaussian fits to the extension histograms. As a check for consistency, $\Delta x$ was also determined by partitioning extension into two or more states using the Hidden Markov Chain analysis similar to that previously described (2). This value was found to agree (within experimental uncertainties) with the value obtained from a direct Gaussian fit.

The number of RNA nucleotides involved in each transition was calculated from the extension change ($\Delta x$) at a given force by adding or subtracting the width of an A-form helix and dividing by the extension expected per nucleotide ($l_F$) at that force (Equation 3), assuming a modified Marko-Siggia worm-like chain (WLC) model (3,4) (Equation 4) for ssRNA.

$$\Delta N_{ssRNA} = (\Delta x + D \cdot \Delta N_{\text{helix}})/l_F \tag{3}$$

$$F(x) = \frac{k_B T}{P}\left[\frac{1}{4}\left(1 - \frac{x}{L_c}\right)^{-2} - \frac{1}{4} + \frac{x}{L_c} - \frac{F}{S}\right] \tag{4}$$

Here $k_B$ is the Boltzmann constant, $T$ is the absolute temperature (297 K), $D$ is the diameter of A-form dsRNA helix (2.2 nm), $L_c$ is the contour length, $P$ is the persistence length and $S$ is the stretch modulus. For ssRNA, $L_c$ is 0.59 nm per nucleotide, $P$ is 1 nm and $S$ is 1500 pN, respectively (5). At each force, the value of $l_F$ can be calculated according to Equation 4. Once the numbers of unfolded helices ($\Delta N_{helix}$) were determined, the numbers of RNA nucleotide changed between two states ($\Delta N_{ssRNA}$) could be obtained.

FECs measured under the same buffer conditions were aligned to remove residual instrumental drift as well as measurement uncertainty of contour length for different tethers, typically a few nanometers. The align method was described previously (6). The force of FECs was aligned using the low force regions (~1 pN) where the force changed very slowly with extension to correct the drifting in force signal. The extension of FECs was aligned using the high force regions (~20 pN) where the force changed very rapidly with extension to correct the drifting in extension. Contour length change of the fully unfolded state was found by fitting the fully unfolded state to an extensible worm-like chain (WLC) model that consisted of two WLCs in series: one for the hybrid handles, and the other for ssRNA that was unfolded in each state. Parameters for handle were first determined by fitting the FECs for the fully folded state. Then the FECs for the fully unfolded states were fitted by treating *P* and *S* as fixed variables, as described above, and thus the contour length of unfolded ssRNA was the only fitting parameter for the fully folded states. Using this model, the nucleotides changed from the fully folded state to the fully unfolded state were fitted. However, the intermediate states in the FECs tend to transit rapidly, leaving large uncertainty, which was difficult to fit. Thus, the WLC curves for the intermediate states '$I_1$', '$I_2$', '$I_3$' and '$I_4$' were plot according to this sequential WLC model using ideal contour lengths estimated from force-clamp experiments.

*Determination of force-dependent kinetic rates*

For force-clamp experiments, the force-dependent kinetic rates were directly obtained by applying Hidden Markov Chain analysis to the extension trajectories using the ebFRET MATLAB program developed by van de Meent et al (2). Since folding of the tRNA was apparently sequential, each transition in force-clamp experiments could be regarded as a two-state system. We assume the positions of the barriers are force-independent, so that for each transition, the distance of the transition state from the folded or unfolded state along the reaction coordinate, $\Delta x^{\ddagger}$, was determined by (7)

$$k(F) = k_0 \exp\left(F\Delta x^{\ddagger}/k_B T\right) \quad (5)$$

Here $k_B$ is Boltzmann's constant, $k_0$ is the apparent folding/unfolding rate at 0 pN, and $T$ is the absolute temperature. This method was used for the transition state distances from both the folded and unfolded states, based on their respective folded and unfolded state kinetics. The activation energy under a certain force for the two-state kinetic, is determined by (6)

$$\Delta G^{\ddagger}(F) = -k_B T ln\left(\frac{k_{1/2}}{k_p}\right) + (F_{1/2} - F) \cdot \Delta x^{\ddagger} \quad (6)$$

Here $F_{1/2}$, the critical force (where the molecule spends equal time in folded and unfolded states), was determined from the force at which the folding and unfolding rates were equal, i.e., to $k_{1/2}$, and the prefactor $k_p$ is $10^5$ $s^{-1}$, determined by Woodside *et al* (8). Due to the experimental uncertainties, the $\Delta x^{\ddagger}$ used in equation 6 here was scaled so that the sum of absolute values of $\Delta x^{\ddagger}$ from both the folded and unfolded states equals to the total $\Delta x$ of the unfolding step, as described by Greenleaf *et al* (9).

For the pulling experiments, the force-dependent unfolding rate and location of the barrier for the transition $F \rightarrow I_1$ are determined by analyzing the distributions of the unfolding forces from the FECs. An analytical expression proposed by Dudko *et al* (10,11) was used to fit the distributions of rupture forces (Figure 5A-C, insets) to parameterize the initial energy barrier to unfolding in terms of $\Delta G_0^{\ddagger}$ (the activation energy at zero force), $\Delta x^{\ddagger}$ (the distance to the barrier at zero force), and $k_{off}$ (the intrinsic off-rate).

$$p(\mathrm{F}) \propto \frac{k(F)}{r} \exp\left\{\frac{k_{\mathrm{off}}}{\Delta x^{\ddagger} r} - \frac{k(F)}{\Delta x^{\ddagger} r}\left(1 - \frac{\Delta x^{\ddagger} F}{\Delta G_0^{\ddagger}}\right)^{1-\frac{1}{v}}\right\} \tag{7}$$

where $k(F) = k_{off}(1 - \frac{\Delta x^{\ddagger} F}{\Delta G_0^{\ddagger}})^{\frac{1}{v}-1} exp\left\{\frac{\Delta G_0^{\ddagger}}{k_B T}\left[1 - (1 - \frac{\Delta x^{\ddagger} F}{\Delta G_0^{\ddagger}} v)^{1/v}\right]\right\}$

Here, $r$ is the force loading rate, and the input parameter, $v$, was alternatively set to 1/2 or 2/3, corresponding to a cusp-like or linear-cubic potential, respectively. Since the shape of the potential is unknown, we averaged the fitting results by setting $v$=1/2 and $v$=2/3. As the $\Delta x^{\ddagger}$ is close to the 'F' state, we assume that the barrier for the transition F→$I_1$ is rigid, so that its position is force-independent. The force-dependent activation energy was then calculated (6):

$$\Delta G^{\ddagger}(F) = \Delta G_0^{\ddagger} - F \cdot \Delta x^{\ddagger} \tag{8}$$

*Determination of free energies*

For the force-clamp experiments, the free energy difference between adjacent states within the landscape was calculated from (7)

$$\Delta G(F) = (F_{1/2} - F) \cdot \Delta x \tag{9}$$

The energy landscapes include the free energy to unfold ssRNA during the certain transition to the force *F*. The force-clamp results used to generate the energy landscapes are listed in Supplementary Tables S1, S2, S5, S6. All energies and positions in the landscape were calculated with reference to the state 'U'.

For the pulling experiments, the free energy difference between 'F' and '$I_1$' states at zero force was determined by Jarzynski equality (12). Firstly, the work required to unfold the whole structure, including handles, was calculated by integrating FECs of the F→$I_1$. Then, the energy required to stretch out the handles and ssRNA to the identical force must be subtracted from this work. The latter energy is calculated by integrating the FEC expected for ssRNA part of '$I_1$' out to the force where the F→$I_1$ transition ended. The Jarzynski estimator is known to have a systematic bias when only finite numbers of measurements are sampled, due to nonlinear weighting of the data. We corrected the bias using the method described by Jeff Gore *et al* previously (13), as 4 ± 1 kcal/mol. The force-dependent free energy between F and $I_1$ was then calculated (7):

$$\Delta G(F) = \Delta G_{0,JE} + \Delta G_{ssRNA} - F \cdot \Delta x \tag{10}$$

Here $\Delta G_{0,JE}$ is the free energy difference at 0 force determined by Jarzynski equality as described above, and $\Delta G_{ssRNA}$ is the free energy to stretch the ssRNA part of '$I_1$', calculated by integrating Equation 4 from 0 to the extension at force *F*.

**Assignment of folding states of the wildtype tRNA$^{phe}$ in the pulling experiment**

To further assign the folding states in the single-molecule pulling experiments, we first fitted the overlapped FEC data of different states to a model with two extensible worm-like chains (EWLC) in series (Equation 4, see Main text, material and methods), to determine the total increased number of $\Delta N_{ssRNA}$ due to tRNA unfolding (Main Text, Figure 2D, solid lines) (6). For the fully unfolded state ('U'), where $\Delta N_{helix}$ = -1, we observed that $\Delta N_{ssRNA}$ = 71.5 ± 0.7 nt (mean ± S.E.,

N = 75), in good agreement with the total length (72 nt) of the tRNA (excluding the 3' ACCA tail). Next, for the intermediate states, we used the $\Delta N_{ssRNA}$ values from the force-clamp experiments (Equation 4, see Main text, material and methods) to calculate the theoretical traces (Main Text, Figure 2D, dashed lines) and compared them with the measured FECs. Immediately, we noticed that the intermediate state in the two-step unfolding pathway is well aligned with the theoretical curve of '$I_1$', indicating that '$I_1$' is the intermediate state in the unfolding process. The first (large) force rupture observed in the unfolding process thus corresponds to the disruption of the major tertiary interactions of tRNA as well as other secondary structures (such as AS, TSL, DSL, etc.) except ASL all at once. The second (small) force rupture represents further disruption of ASL to completely unfold tRNA. For the refolding pathway in tRNA pulling experiments, the first refolding intermediate state was also well aligned with theoretical curve of '$I_1$' as shown in Figure 2D. The intermediate '$I_3$' and '$I_4$' were recognized in the refolding pathway as well. As the transitions between the refolding intermediates merge in the FECs, it was difficult to recognize the state '$I_2$'.

**The possible structure of '$I_3$' state**

The '$I_3$' state may represent a structure comprising three stem-loops, while the VL and the AS regions are still single-stranded under tension. According to this assumption, the value of experimental $\Delta N_{ssRNA}$ for the $I_3 \rightarrow F$ transition was calculated using Equation 3 (Supplementary Table S1, S5), which equals to 24.6 ± 0.8 nt (wildtype) and 23.8 ± 0.5 nt (mutant) (assuming $\Delta N_{helix}$ = -2, as the formation of the AS reduced the apparent number of folded stems by two). The experimental $\Delta N_{ssRNA}$ value for the $I_3 \rightarrow F$ transition agrees with the theoretical $\Delta N_{ssRNA}$ value of 22 nt. The $I_3 \leftrightarrow I_4$ transition corresponds to the coaxial stacking of ASL and DSL ($\Delta N_{helix}$ = -1), leading to the folding of G26 as well as A44, G45 and G46 in the VL (so that theoretical $\Delta N_{ssRNA}$ = 4 nt). The experimental $\Delta N_{ssRNA}$ value for $I_3 \leftrightarrow I_4$ transition also agrees with the theoretical value (Supplementary Table S1, S5).

Alternatively, '$I_3$' may represent a structure in which the TSL is not folded but ASL and DSL are coaxially stacked and G26 as well as A44, G45 and G46 in the VL are folded. In this case, for the $I_2 \leftrightarrow I_3$ transition, the $\Delta N_{helix}$ = -1 and the theoretical $\Delta N_{ssRNA}$ = 4; for the $I_3 \leftrightarrow I_4$ transition, the $\Delta N_{helix}$ = 1 and the theoretical $\Delta N_{ssRNA}$ = 17 nt (corresponding to the folding of TSL), and for $I_3 \rightarrow F$ transition, the $\Delta N_{helix}$ = 0 and the theoretical $\Delta N_{ssRNA}$ = 35 nt (corresponding to the folding of U8, A9, U47, C48, AS and TSL). Based on this assumption, we calculated the experimental $\Delta N_{ssRNA}$ values of wildtype and mutant tRNA for transitions $I_2 \leftrightarrow I_3$, $I_3 \leftrightarrow I_4$ and $I_3 \rightarrow F$/ $I_3 \leftrightarrow F^*$ (Supplementary Table S1, S5). All the experimental $\Delta N_{ssRNA}$ values are consistent with the theoretical ones, thus both assumptions can explain the conformation of '$I_3$'.

**The possible structure of '$I_4$' and '$I_4$'-like state**

The '$I_4$'-like state occurred at low salt condition could be explained by the salt-dependent stability of the AS due to the G4◦U69 wobble base pair, which has strong negative potential in its major groove (14). Such negative potential destabilizes the acceptor stem when the ionic strength of the buffer is low. We noticed that '$I_4$' presented in the folding pathways of both the wildtype tRNA and the mutant, suggesting that the elbow base pairings (G18-U55 and G19-C56) between ASL and DSL as well as the Levitt pair between DSL may not take part in the formation of '$I_4$'. Moreover, '$I_4$' is unlikely to be a misfolded state, as it is known that the topological constraints of the junction of tRNA suppress the formation of misfolded intermediates (15). Therefore, '$I_4$' probably represents a structure that the D stem, anti-codon stem and variable loop stack coaxially by formation of base triples C13-G22◦G46, G10-C25◦G45 and non-canonical base pair G26◦A44, while U8◦A14, U12-A23◦A9, G15◦C48 and the acceptor stem do not form. We did not observe $I_4$→F transition during all the force-clamp experiments, probably because the $I_4$→F transition is very slow at the force range we performed force-clamp experiments.

**Supplementary Figures and Tables**

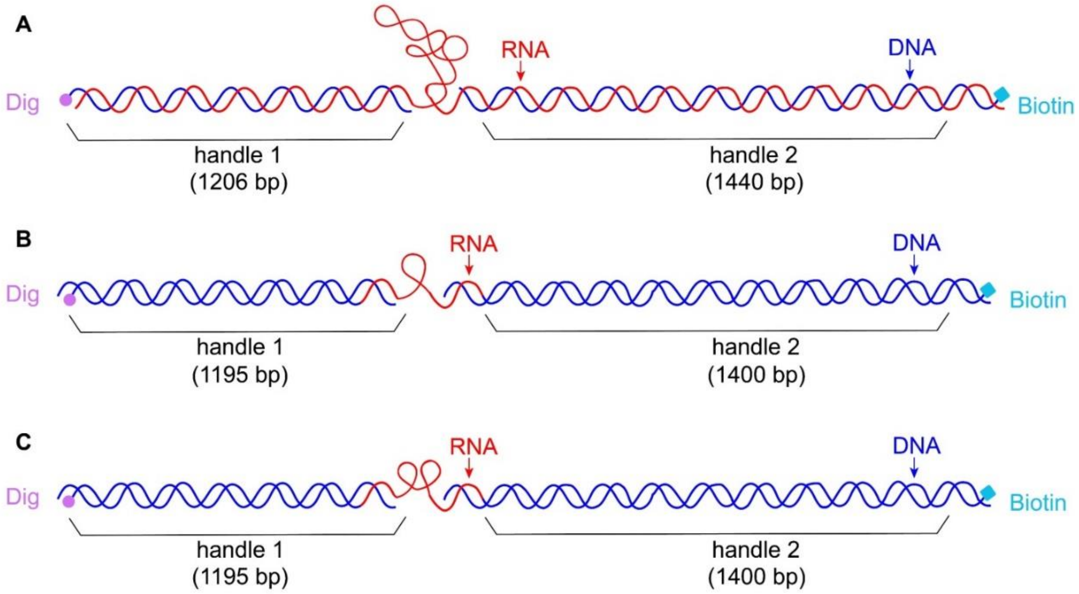


**Supplementary Figure S1.** Single-molecule constructs of the samples. (A) Wildtype and mutant yeast tRNA$^{phe}$ samples. (B) Individual ASL, DSL and TSL samples. (C) Truncated tRNA constructs DAV and AVT samples.

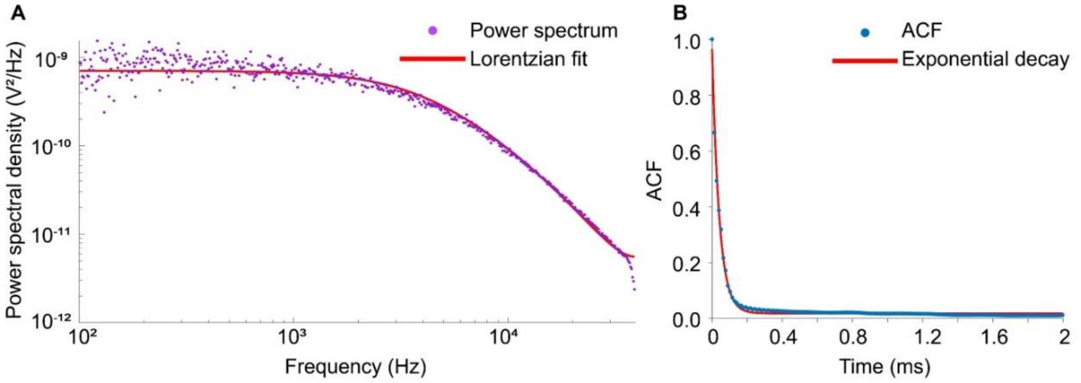


**Supplementary Figure S2.** A typical power spectrum and autocorrelation function (ACF) measured in a commercial dual-trap optical tweezers (LUMICKS, C-Trap). (A) The power spectrum density recorded at 78.125 kHz sampling frequency to determine the trap stiffness, the cut-off frequencies ($f_c$) was fit using Lorentzian function (16), yielding $f_c$ = 4127 Hz. (B) The autocorrelation function (ACF) was measured by stretching a 2.8 kbp control DNA at a constant force (8 pN) and recording the force signal at 78.125 kHz. The fitting result indicates that the $\tau_0$ = 0.040 ms.

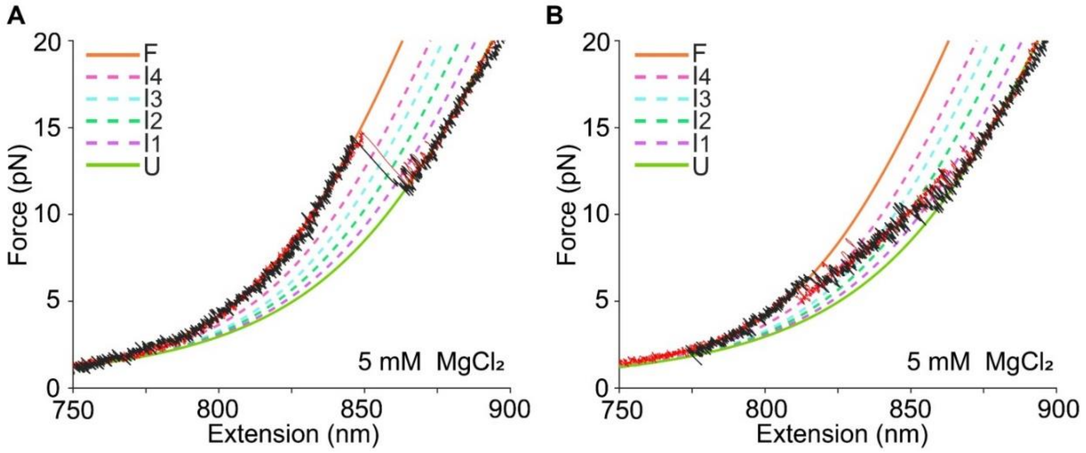


**Supplementary Figure S3.** Comparison of force extension curves (FEC) acquired by single-trap optical tweezers and dual-trap optical tweezers. (A) Typical unfolding FEC of wildtype tRNA$^{phe}$ in 5mM $MgCl_2$ acquired by single-trap optical tweezers (black) and dual-trap optical tweezers (red). (B) Typical refolding FEC of wildtype tRNA$^{phe}$ in 5 mM $MgCl_2$ acquired by single-trap optical tweezers (black) and dual-trap optical tweezers (red).

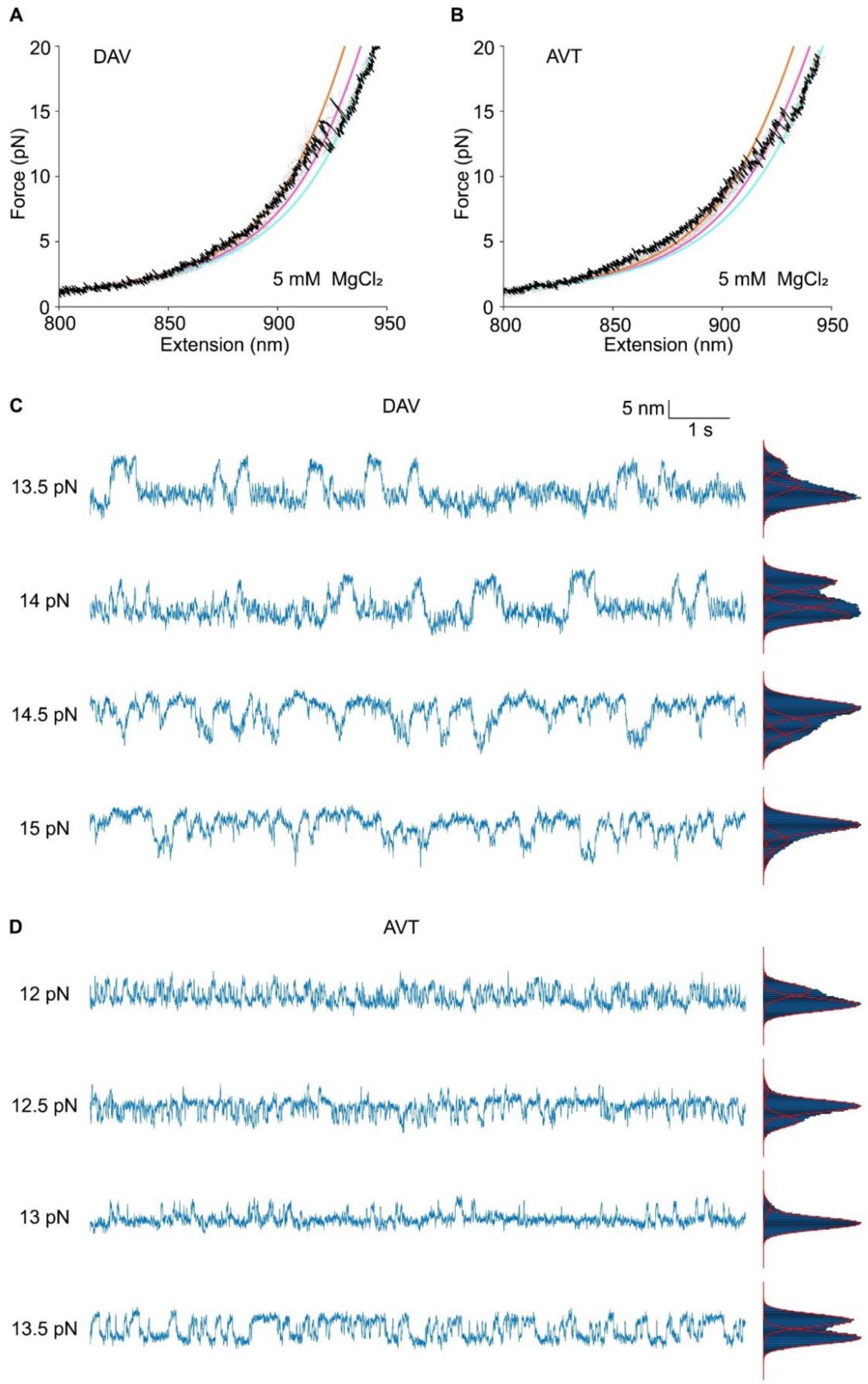


**Supplementary Figure S4.** Force measurement of two truncated tRNA constructs DAV and AVT. Black curves represent typical unfolding trajectories of DAV (A) and AVT (B) are plotted above the aggregated data (grey dots). Typical unfolding trajectories of DAV (C) and AVT (D) acquired in force clamp experiment were showed. Data were smoothed to 200 Hz. The extension histogram was fitted to Gaussian functions.

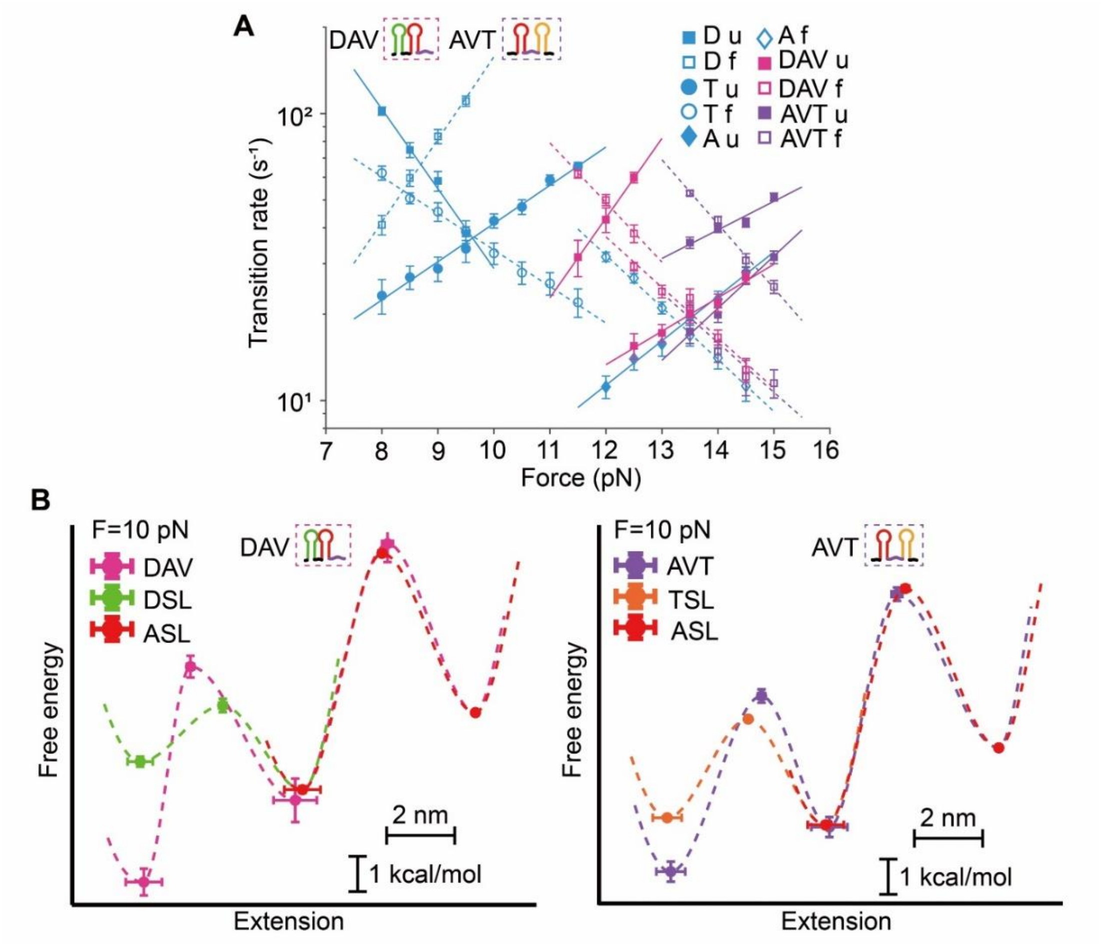


**Supplementary Figure S5.** Folding and unfolding details of truncated tRNA constructs DAV and AVT. (A) Force-dependent kinetics of unfolding/folding of each transition of truncated tRNA constructs DAV and AVT as well as individual ASL, DSL and TSL during force-clamp experiments. (B) Free energy landscapes of truncated tRNA constructs DAV and AVT as well as individual ASL, DSL and TSL under 10 pN at 5 mM $MgCl_2$. The free energy landscapes of DAV, AVT and individual ASL are plotted with reference to the unfolded state, and the free energy landscapes of individual DSL and TSL are plotted with reference to the folded state of individual ASL. Error bars showed as S.E.

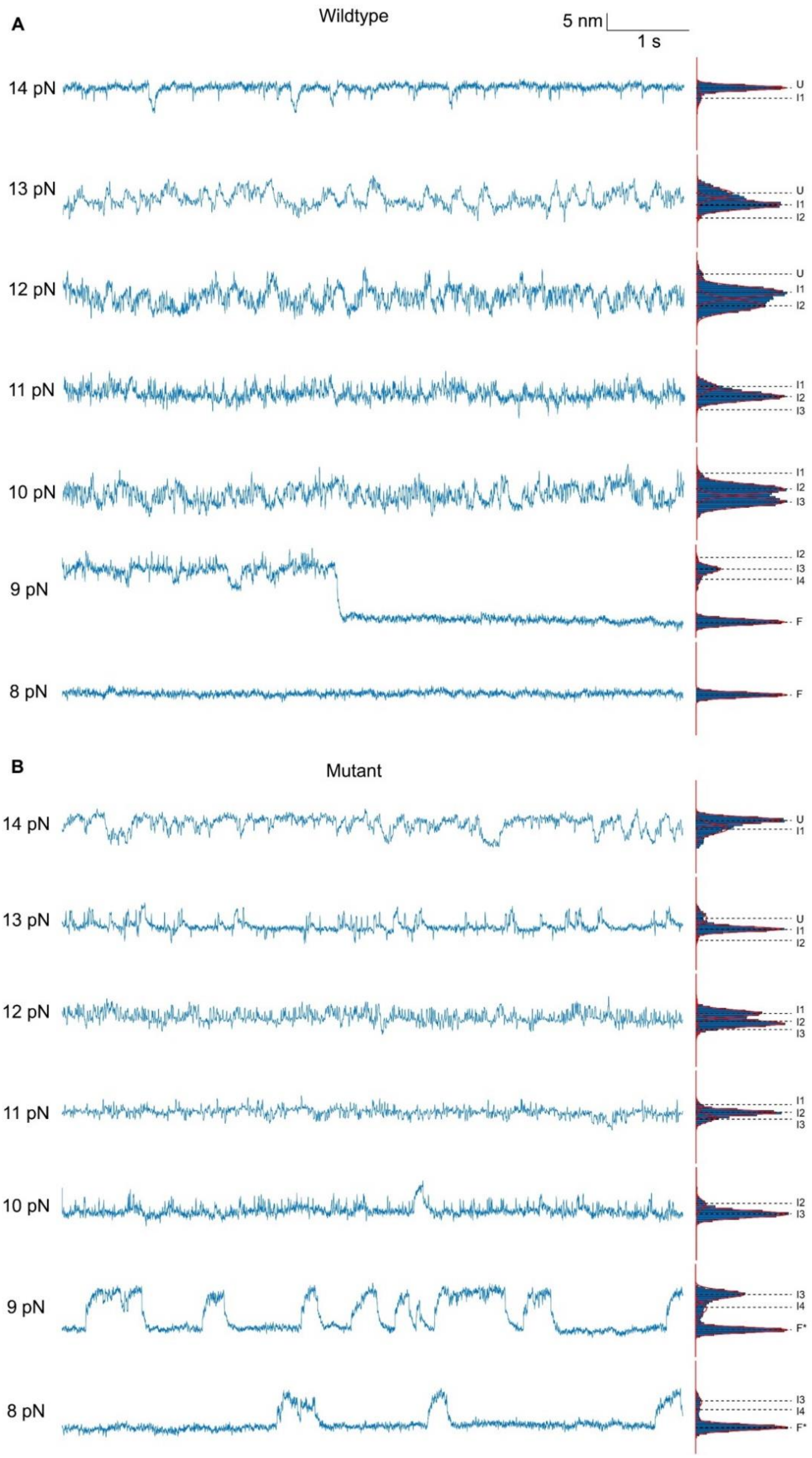


**Supplementary Figure S6.** Stepwise refolding of (A) wildtype and (B) mutated tRNA$^{phe}$. Data were smoothed to 200 Hz. The extension histogram was fitted to Gaussian functions.

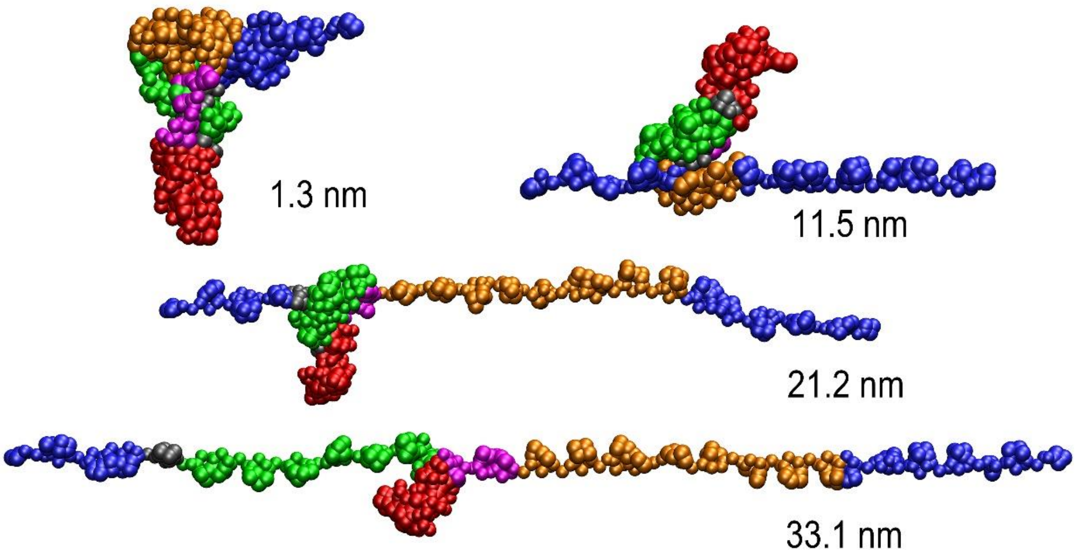


**Supplementary Figure S7.** Snapshots of the representative conformations of the wildtype $tRNA^{phe}$ during unfolding in the presence of $Mg^{2+}$ at the pulling rate of 0.2 nm/ns from CG SMD simulations. The extension corresponding to each conformation is shown, respectively. CG particles are shown as spheres colored to identify the different regions of secondary structure (AS, blue; TSL, yellow; DSL, green; ASL, red; VL, purple).

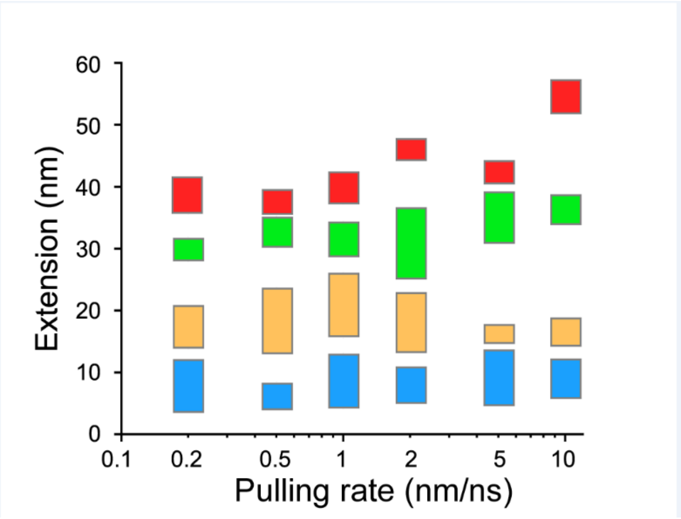


**Supplementary Figure S8.** The effect of loading rate on the unfolding sequence of the secondary structures of tRNA in CG SMD simulations. The bars of extension spans represent the onset and the end of stem-loop disruptions (AS, blue; TSL, yellow; DSL, green; ASL, red).

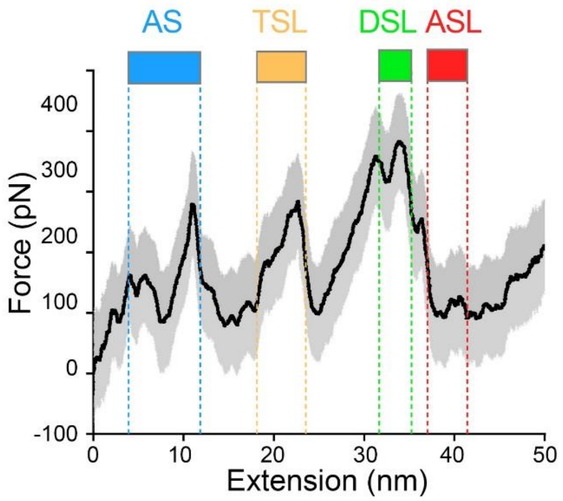


**Supplementary Figure S9.** Representative force-extension curves by CG SMD simulations initiating from the 'F' state without cations. The black lines represent the running averages with a window size of 400 ps. The gray shades denote the standard deviations of the running averaging. Colored bars and dashed lines represent the extension spans indicating the onset and the end of stem-loop disruptions (AS, blue; TSL, yellow; DSL, green; ASL, red).

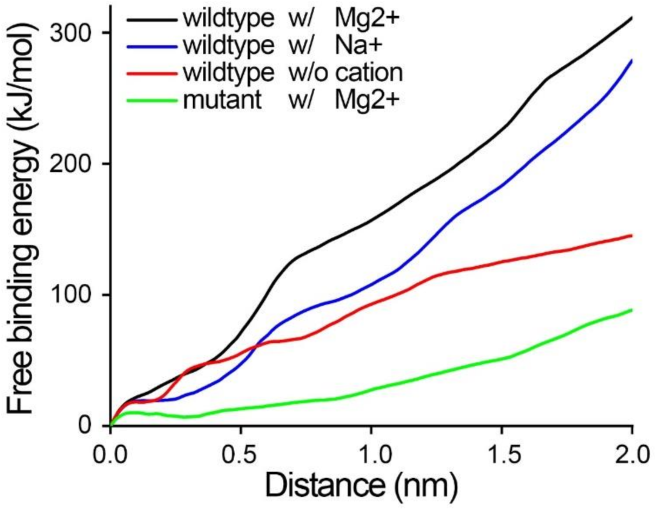


**Supplementary Figure S10.** Free binding energy profile of tRNA transforming from the canonical L-shape folded state (‘F’) to the elbow-disrupted folded state (‘F*’) at different cation conditions from all-atom SMD simulations. Black, blue, red and green lines represent the free binding energies of the wildtype tRNA with $Mg^{2+}$, the wildtype tRNA with $Na^{+}$, the wildtype tRNA without cations and the mutant tRNA with $Mg^{2+}$, respectively.

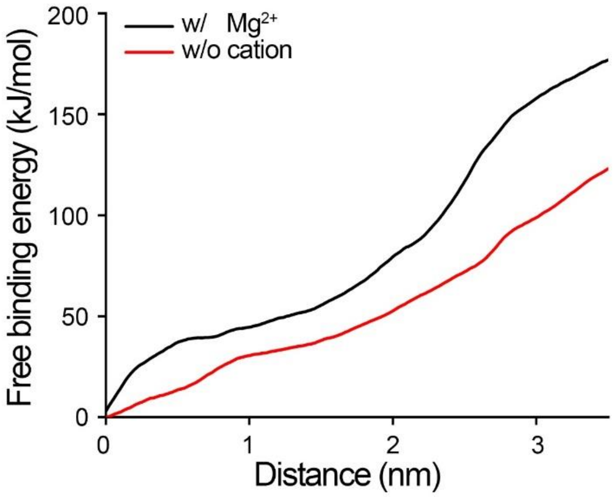


**Supplementary Figure S11.** Free binding energy profile of tRNA transforming from the canonical L-shape folded state (‘F’) to the AS-disrupted state (‘$I_4$’) at different cation conditions from all-atom SMD simulations. Black and red lines represent the free binding energies of the wildtype tRNA with $Mg^{2+}$ and wildtype tRNA without cations, respectively.

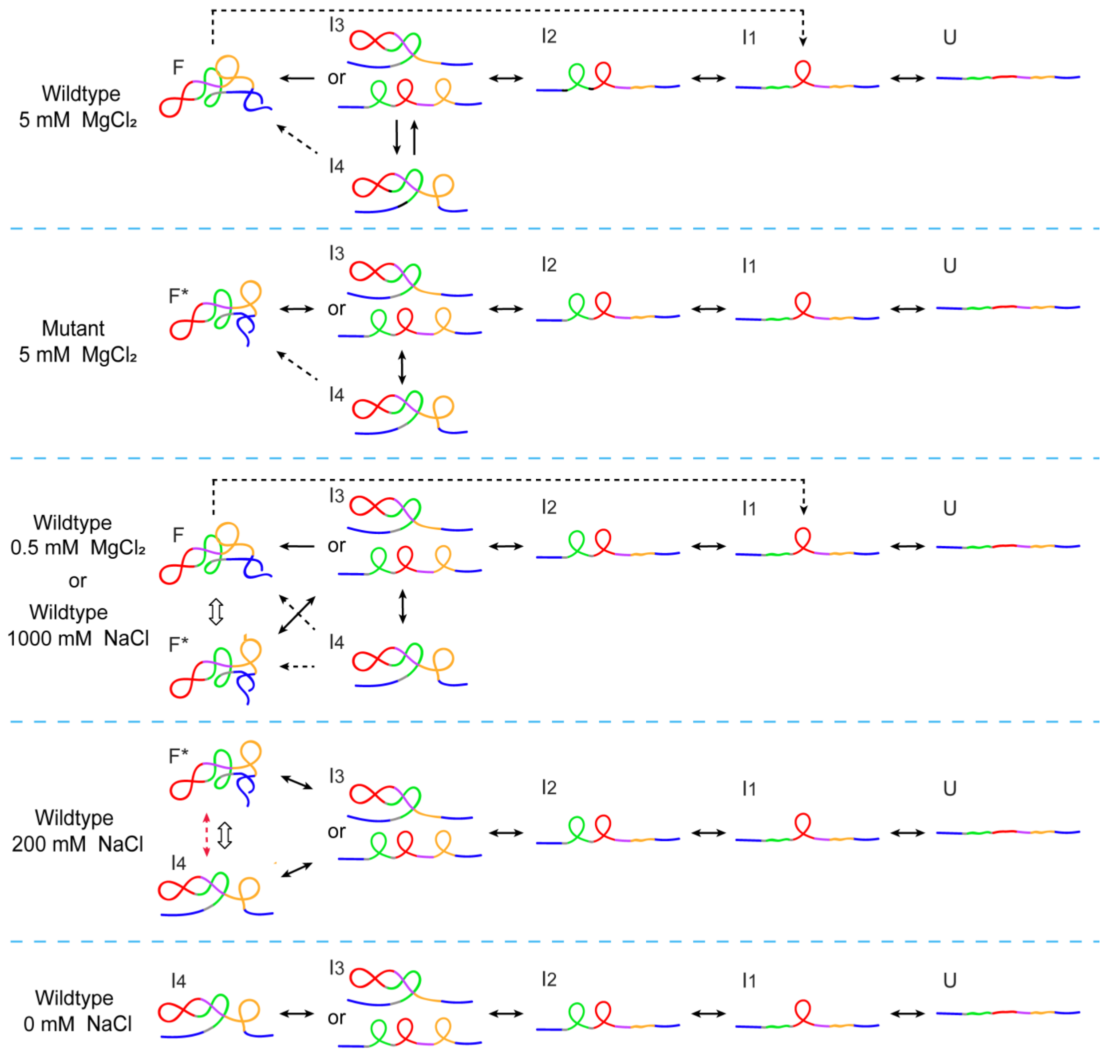


**Supplementary Figure S12.** Summary of the folding and unfolding pathways of tRNA$^{phe}$ under different circumstances. Solid arrows indicate transitions observed in force-clamp experiments, dashed arrows indicate transitions observed in pulling experiments, and hollow arrows indicate transitions at low force which we did not directly observed.

**Supplementary Movie S1 and Movie S2 (separate file).** Steered molecular dynamics simulation results of elbow-formed (Movie 1) and elbow-disrupted (Movie 2) tRNA unfolding. Each movie contains 2225 frames and the simulation time interval is 100 ps/frame, with a pulling rate of 0.2 nm/ns. The final stretched tRNA molecule is 44.5 nm in length.

**Supplementary Table S1. Transitions of wild type $tRNA^{phe}$ in the constant force experiment at 5 mM $MgCl_2$.** The errors are presented as mean ± S.E. from 10 traces.

| Transition | $\Delta x$ (nm) | Theoretical $\Delta N_{ssRNA}$ (nt) | Experimental $\Delta N_{ssRNA}$ (nt) | $\Delta N_{helix}$ |
|---|---|---|---|---|
| $I_3 \to F$ | 13.5 ± 0.2 | 22[a] | 24.6 ± 0.8[a] | -2[a] |
| | | 35[b] | 36.5 ± 0.8[b] | 0[b] |
| $I_3 \leftrightarrow I_4$ | 3.9 ± 0.7 | 4[a] | 4.6 ± 1.8[a] | -1[a] |
| | | 17[b] | 16.5 ± 1.8 | 1[b] |
| $I_2 \leftrightarrow I_3$ | 4.3 ± 0.5 | 17[a] | 16.2 ± 1.3[a] | 1[a] |
| | | 4[b] | 5.2 ± 1.3[b] | -1[b] |
| $I_1 \leftrightarrow I_2$ | 4.9 ± 0.6 | 16 | 17.1 ± 1.5 | 1 |
| $U \leftrightarrow I_1$ | 4.4 ± 0.4 | 17 | 15.7 ± 1.0 | 1 |

[a] Assuming that $I_3$ corresponds to a conformation with ASL, DSL and TSL folded but no inter-helical stacking.

[b] Assuming that $I_3$ corresponds to a conformation with ASL and DSL coaxial stacked and the G26 as well as A44, G45 and G46 in the VL folded but the TSL is not folded.

**Supplementary Table S2. Unfolding and folding kinetics of wild type $tRNA^{phe}$ in the constant force experiment at 5mM $MgCl_2$.** Data are presented as mean ± S.E. from 10 traces.

| Transition | $\ln k_0$ ($s^{-1}$) | $x^‡$ (nm) | $F_{1/2}$ (pN) | $\ln(k_{1/2})$ ($s^{-1}$) | $\Delta G^‡_{1/2}$ (kcal/mol) |
|---|---|---|---|---|---|
| $I_3 \to I_2$ | 0.8 ± 0.1 | 1.3 ± 0.1 | 10.1 ± 0.2 | 4.5 ± 0.1 | 2.8 ± 0.1 |
| $I_2 \to I_3$ | 12.5 ± 0.4 | -3.3 ± 0.2 | | | |
| $I_2 \to I_1$ | 3.3 ± 0.1 | 0.5 ± 0.1 | 10.4 ± 0.2 | 4.2 ± 0.1 | 3.0 ± 0.1 |
| $I_1 \to I_2$ | 11.4 ± 0.2 | -2.8 ± 0.1 | | | |
| $I_1 \to U$ | -2.0 ± 0.3 | 1.5 ± 0.1 | 13.5 ± 0.6 | 2.9 ± 0.2 | 3.7 ± 0.1 |
| $U \to I_1$ | 8.4 ± 0.4 | -1.7 ± 0.1 | | | |

**Supplementary Table S3. Transitions of individual stem-loops in the constant force experiment at 5 mM $MgCl_2$.** The transition distance $\Delta x$ is presented as mean ± S.E., as determined by Gaussian fitting of extension histogram at force near $F_{1/2}$.

| Transition | $\Delta x$ (nm) | $\Delta N_{ssRNA}$ (nt) | $\Delta N_{helix}$ |
|---|---|---|---|
| DSL↔U | 4.5 ± 0.4 | 15.6 ± 1.1 | -1 |
| TSL↔U | 4.4 ± 0.4 | 16.0 ± 1.1 | -1 |
| ASL↔U | 4.8 ± 0.5 | 16.0 ± 1.2 | -1 |

**Supplementary Table S4. Unfolding and folding kinetics of individual stem-loops in the constant force experiment at 5 mM $MgCl_2$.** Folding and unfolding kinetics were recorded by holding the tether at each force for > 40 s. Data are presented as mean ± S.E. from 14 traces (DSL), 10 traces (TSL) and 12 traces (ASL), respectively. Typically, 1-3 traces were measured from an individual molecule in force clamp experiment.

| Transition | ln$k_0$ ($s^{-1}$) | $x^‡$ (nm) | $F_{1/2}$ (pN) | ln($k_{1/2}$) ($s^{-1}$) | $ΔG^‡_{1/2}$ (kcal/mol) |
|---|---|---|---|---|---|
| DSL→U | -1.5 ± 0.3 | 2.7 ± 0.2 | 8.7 ± 0.5 | 4.2 ± 0.3 | 2.9 ± 0.2 |
| U→DSL | 9.8 ± 0.5 | -2.6 ± 0.3 | | | |
| TSL→U | 0.7 ± 0.1 | 1.3 ± 0.1 | 9.6 ± 0.2 | 3.6 ± 0.1 | 3.3 ± 0.1 |
| U→TSL | 6.5 ± 0.1 | -1.2 ± 0.1 | | | |
| ASL→U | -1.9 ± 0.2 | 1.5 ± 0.1 | 13.3 ± 0.3 | 2.9 ± 0.1 | 3.7 ± 0.1 |
| U→ASL | 8.1 ± 0.1 | -1.7± 0.1 | | | |

**Supplementary Table S5. Transitions of the mutant tRNA in the constant force experiment at 5 mM $MgCl_2$.** The transition distance $Δx$ is presented as mean ± S.E., while other parameters are presented as mean ± S.E. from 12 traces.

| Transition | $Δx$ (nm) | Theoretical $ΔN_{ssRNA}$ (nt) | Experimental $ΔN_{ssRNA}$ (nt) | $ΔN_{helix}$ |
|---|---|---|---|---|
| $I_3$↔$F^*$ | 13.2 ± 0.2 | 22[a] | 23.8 ± 0.5[a] | -2[a] |
| | | 35[b] | 35.7 ± 0.5[b] | 0[b] |
| $I_3$↔$I_4$ | 4.1 ± 0.4 | 4[a] | 5.1 ± 1.1[a] | -1[a] |
| | | 17[b] | 16.9 ± 1.1[b] | 1[b] |
| $I_2$↔$I_3$ | 4.5 ± 0.3 | 17[a] | 16.7 ± 0.8[a] | 1[a] |
| | | 4[b] | 5.7 ± 0.8[b] | -1[b] |
| $I_1$↔$I_2$ | 4.8 ± 0.5 | 16 | 17.1 ± 1.2 | 1 |
| U↔$I_1$ | 4.5 ± 0.4 | 17 | 16.0 ± 1.0 | 1 |

[a] Assuming that $I_3$ corresponds to a conformation with ASL, DSL and TSL folded but no inter-helical stacking.

[b] Assuming that $I_3$ corresponds to a conformation with ASL and DSL coaxial stacked and the G26 as well as A44, G45 and G46 in the VL folded but the TSL is not folded.

**Supplementary Table S6. Unfolding and folding kinetics of the mutant tRNA in the constant force experiment at 5 mM $MgCl_2$.** Data are presented as mean ± S.E. from 12 traces.

| Transition | ln$k_0$ ($s^{-1}$) | $x^‡$ (nm) | $F_{1/2}$ (pN) | ln($k_{1/2}$) ($s^{-1}$) | $ΔG^‡_{1/2}$ (kcal/mol) |
|---|---|---|---|---|---|
| $F^*$→$I_3$ | -9.4 ± 0.2 | 4.2 ± 0.1 | 9.5 ± 1 | 0.3 ± 0.1 | 5.2 ± 0.5 |
| $I_3$→$F^*$ | 13.1 ± 1.8 | -5.5 ± 0.7 | | | |
| $I_3$→$I_2$ | -1.5 ± 0.7 | 2.6 ± 0.3 | 9.6 ± 0.3 | 4.6 ± 0.3 | 2.7 ± 0.1 |
| $I_2$→$I_3$ | 12.2 ± 0.4 | -3.3 ± 0.2 | | | |
| $I_2$→$I_1$ | -2.8 ± 0.3 | 2.5 ± 0.1 | 12.1 ± 0.1 | 4.8 ± 0.2 | 2.6 ± 0.1 |
| $I_1$→$I_2$ | 13.3 ± 0.6 | -2.9 ± 0.2 | | | |
| $I_1$→U | -1.4 ± 0.6 | 1.3 ± 0.2 | 13.3 ± 0.9 | 2.9 ± 0.4 | 3.7 ± 0.2 |
| U→$I_1$ | 8.5 ± 0.4 | -1.7 ± 0.1 | | | |

**Supplementary Table S7. Unfolding rate and transition position of the transition F→$I_1$ in the presence of $MgCl_2$.** Data are presented as mean ± S.E. from 65 traces (0.5 mM $MgCl_2$), 58 traces (2 mM $MgCl_2$) and 75 traces (5 mM $MgCl_2$), respectively. Typically, 3-5 traces were measured from an individual molecule in pulling experiment.

| $MgCl_2$ conc. | $k_0$ ($s^{-1}$) | $x^‡$ (nm) | $ΔG^‡$ (kcal/mol) |
|---|---|---|---|
| 0.5 mM | $(9.2 ± 0.2)×10^{-8}$ | 6.8 ± 0.4 | 21.3 ± 2.6 |
| 2 mM | $(1.5 ± 0.1)×10^{-7}$ | 6.2 ± 0.3 | 19.3 ± 1.5 |
| 5 mM | $(1.2 ± 0.1)×10^{-7}$ | 6.1 ± 0.4 | 16.3 ± 0.4 |

**Supplementary Table S8. Transitions of DAV in the constant force experiment at 5 mM $MgCl_2$.** The errors are presented as mean ± S.E. from 11 traces.

| Transition | $Δx$ (nm) | $ΔNssRNA$ (nt) | $ΔN_{helix}$ |
|---|---|---|---|
| ASL↔F | 4.2± 0.5 | 14.6 ± 1.2 | -1 |
| U↔ASL | 5.0 ± 0.6 | 16.7 ± 1.4 | -1 |

**Supplementary Table S9. Unfolding and folding kinetics of DAV in the constant force experiment at 5 mM $MgCl_2$.** Data are presented as mean ± S.E. from 11 traces.

| Transition | ln$k_0$ ($s^{-1}$) | $x^‡$ (nm) | $F_{1/2}$ (pN) | ln($k_{1/2}$) ($s^{-1}$) | $ΔG^‡_{1/2}$ (kcal/mol) |
|---|---|---|---|---|---|
| F→ASL | 0.5 ± 0.8 | 0.9 ± 0.2 | 14.1 ± 0.2 | 3.7 ± 0.1 | 3.2 ± 0.1 |
| ASL→F | 11.0 ± 0.6 | -2.1 ± 0.2 | | | |
| ASL→U | -2.8 ± 0.8 | 1.7 ± 0.2 | 13.6 ± 1.2 | 2.9 ± 0.9 | 3.7 ± 0.5 |
| U→ASL | 8.3 ± 1.7 | -1.6 ± 0.5 | | | |

**Supplementary Table S10. Transitions of AVT in the constant force experiment at 5 mM $MgCl_2$.** Data are presented as mean ± S.E. from 12 traces.

| Transition | $\Delta x$ (nm) | $\Delta NssRNA$ (nt) | $\Delta N_{helix}$ |
|---|---|---|---|
| ASL↔F | 4.4 ± 0.4 | 15.6 ± 1.0 | -1 |
| U↔ASL | 4.7 ± 0.5 | 16.0 ± 1.2 | -1 |

**Supplementary Table S11. Unfolding and folding kinetics of AVT in the constant force experiment at 5 mM $MgCl_2$.** Data are presented as mean ± S.E. from 12 traces.

| Transition | $\ln k_0$ ($s^{-1}$) | $x^‡$ (nm) | $F_{1/2}$ (pN) | $\ln(k_{1/2})$ ($s^{-1}$) | $\Delta G^‡_{1/2}$ (kcal/mol) |
|---|---|---|---|---|---|
| F→ASL | -3.9 ± 0.5 | 2.6 ± 0.2 | 12.1 ± 0.9 | 3.8 ± 0.6 | 3.2 ± 0.3 |
| ASL→F | 9.6 ± 0.7 | -2.0 ± 0.3 | | | |
| ASL→U | -0.6 ± 0.3 | 1.1± 0.1 | 13.5 ± 1.0 | 3.0 ± 0.3 | 3.7 ± 0.2 |
| U→ASL | 8.5 ± 0.8 | -1.7± 0.2 | | | |

**Supplementary Table S12. List of DNA oligomers used in sample preparation. All DNA oligomers were purchased from Sangon (Shanghai, China).**

| Name | Sequence (5'to 3') | Notes | Experiments |
|---|---|---|---|
| HAF | GGTGCCTCACTGATTAAGCATTGGTAA | | PCR Primers to synthesis blunt end handle to be labeled with biotin, which was used in the wildtype and mutant tRNA constructs. |
| HAR | AAGCTTGGCGTAATCATGGTCATAGC | | |
| HBF | CTAGAGGATCCCCGGGTACC | | PCR Primers to |

| | | | |
|---|---|---|---|
| Dig-HBR | Dig-TCGTTCATCCATAGTTGCCTGACTC | 5’ end of the oligomer was digoxygenin labeled. | synthesis blunt end handle to be labeled with digoxygenin, which was used in the wildtype and mutant tRNA constructs. |
| T7-HAF | <u>TAATACGACTCACTATA</u>GGTGCCTCACTGATTAAGCATTGGTAA | The underlined sequence indicates the T7 promoter. | PCR Primers to synthesis long DNA templates of wildtype and mutant tRNA constructs for *in vitro* transcription. |
| HBR | TCGTTCATCCATAGTTGCCTGACTC | | |
| Dig-H1F | Dig-GTCGGAACAGGAGAGCGCAC | 5’ end of the oligomer was digoxygenin labeled. | PCR Primers to synthesis digoxigenin labeled auto-sticky handle, which was used in the individual stem-loop and truncated tRNA constructs. |
| H1R | AAGCTTGGCGTAATCATGGTCATAGCTGTT_CCTGTGTGAAATTGTTATCCGCTCACAAT | “_” represents abasic site. | |
| H2F-3S | p-CTAGAGGATCCCCGGGTACCGA | “p” represents 5’ end of the | PCR Primers to synthesis biotin labeled |

| | | | |
|---|---|---|---|
| | GCTCGAATT*C*A*C | oligomer was phosphorylated and "*" represents phosphorothioate bond. | auto-sticky handle, which was used in the individual stem-loop and truncated tRNA constructs. |
| Biotin-H2R | biotin-GAAGCATTTATCAGGGTTATTGTCTCATGAGCGGATACA | 5' end of the oligomer was biotin labeled. | |
| tRNA forward | <u>TAATACGACTCACTATAGG</u>AACAGCTATGACCATGATTACGCCAAG | The underlined sequence indicates the T7 promoter. | DNA oligomers to synthesis short DNA templates for *in vitro* transcription. |
| Anti-TSL | ATTCGAGCTCGGTACCCGGGGATCCTCTAGTCTGTGGATCGAACACAGGAAGCTTGGCGTAATCATGGTCATAGCTGTTCCTATAGTGAGTCGTATTA | | Synthesis of individual TSL construct template for *in vitro* transcription by annealing with oligomer 'tRNA forward'. |
| Anit-ASL | ATTCGAGCTCGGTACCCGGGGATCCTCTAGTCCAGATCTTCAGTCTGGCAAGCTTGGCGTAATCATGGTCATAGCTGTTCCTATAGTGAGTCGTATTA | | Synthesis of individual ASL construct template |

| | | | |
|---|---|---|---|
| | | | for *in vitro* transcription by annealing with oligomer 'tRNA forward'. |
| Anti-DSL | ATTCGAGCTCGGTACCCGGGG ATCCTCTAGCGCTCTCCCAACT GAGCTAAGCTTGGCGTAATCAT GGTCATAGCTGTTCCTATAGTG AGTCGTATTA | | Synthesis of individual DSL construct template for *in vitro* transcription by annealing with oligomer 'tRNA forward'. |
| Anti-DAV | ATTCGAGCTCGGTACCCGGGG ATCCTCTAGGGACCTCCAGATC TTCAGTCTGGCGCTCTCCCAAC TGAGCTAAGCTTGGCGTAATCA TGGTCATAGCTGTTCCTATAGT GAGTCGTATTA | | Synthesis of DAV construct template for *in vitro* transcription by annealing with oligomer 'tRNA forward'. |
| Anti-AVT | ATTCGAGCTCGGTACCCGGGG ATCCTCTAGTCTGTGGATCGAA CACAGGACCTCCAGATCTTCA GTCTGGCAAGCTTGGCGTAATC ATGGTCATAGCTGTTCCTATAG TGAGTCGTATTA | | Synthesis of AVT construct template for *in vitro* transcription by annealing with oligomer 'tRNA forward'. |

### REFERENCES


1. Chandra, V., Hannan, Z., Xu, H. and Mandal, M. (2017) Single-molecule analysis reveals multi-state folding of a guanine riboswitch. *Nat. Chem. Biol.*, **13**, 194-201.
2. van de Meent, J.-W., Bronson, Jonathan E., Wiggins, Chris H. and Gonzalez, Ruben L. (2014) Empirical bayes methods enable advanced population-level analyses of single-molecule FRET experiments. *Biophys. J.*, **106**, 1327-1337.
3. Wang, M.D., Yin, H., Landick, R., Gelles, J. and Block, S.M. (1997) Stretching DNA with optical tweezers. *Biophys. J.*, **72**, 1335-1346.
4. Marko, J.F. and Siggia, E.D. (1995) Stretching DNA. *Macromolecules*, **28**, 8759-8770.
5. Seol, Y., Skinner, G.M. and Visscher, K. (2004) Elastic properties of a single-stranded charged homopolymeric ribonucleotide. *Phys. Rev. Lett.*, **93**, 118102.
6. Neupane, K., Yu, H., Foster, D.A.N., Wang, F. and Woodside, M.T. (2011) Single-molecule force spectroscopy of the add adenine riboswitch relates folding to regulatory mechanism. *Nucleic Acids Res.*, **39**, 7677-7687.
7. Tinoco, I., T. X. Li, P. and Bustamante, C. (2006) Determination of thermodynamics and kinetics of RNA reactions by force. *Q. Rev. Biophys.*, **39**, 325-360.
8. Woodside, M.T., Behnke-Parks, W.M., Larizadeh, K., Travers, K., Herschlag, D. and Block, S.M. (2006) Nanomechanical measurements of the sequence-dependent folding landscapes of single nucleic acid hairpins. *Proc. Natl. Acad. Sci. U.S.A.*, **103**, 6190-6195.
9. Greenleaf, W.J., Frieda, K.L., Foster, D.A.N., Woodside, M.T. and Block, S.M. (2008) Direct observation of hierarchical folding in single riboswitch aptamers. *Science*, **319**, 630-633.
10. Dudko, O.K., Hummer, G. and Szabo, A. (2006) Intrinsic rates and activation free energies from single-molecule pulling experiments. *Phys. Rev. Lett.*, **96**, 108101.
11. Dudko, O.K., Hummer, G. and Szabo, A. (2008) Theory, analysis, and interpretation of single-molecule force spectroscopy experiments. *Proc. Natl. Acad. Sci. U.S.A.*, **105**, 15755-15760.
12. Liphardt, J., Dumont, S., Smith, S.B., Tinoco, I. and Bustamante, C. (2002) Equilibrium information from nonequilibrium measurements in an experimental test of Jarzynski's equality. *Science*, **296**, 1832-1835.
13. Gore, J., Ritort, F. and Bustamante, C. (2003) Bias and error in estimates of equilibrium free-energy differences from nonequilibrium measurements. *Proc. Natl. Acad. Sci. U.S.A.*, **100**, 12564-12569.
14. Varani, G. and McClain, W.H. (2000) The G·U wobble base pair. *EMBO Rep.*, **1**, 18-23.
15. Mustoe, A.M., Brooks, C.L., III and Al-Hashimi, H.M. (2014) Topological constraints are major determinants of tRNA tertiary structure and dynamics and provide basis for tertiary folding cooperativity. *Nucleic Acids Res.*, **42**, 11792-11804.
16. Hansen, P.M., Tolić-Nørrelykke, I.M., Flyvbjerg, H. and Berg-Sørensen, K. (2006) tweezercalib 2.0: Faster version of MatLab package for precise calibration of optical tweezers. *Comput. Phys. Commun.*, **174**, 518-520.
